# The Brugada syndrome associated gene *WT1* impacts on *SCN5A* expression and cardiac conduction

**DOI:** 10.1101/2025.01.16.633330

**Authors:** E. Madelief J. Marsman, Joost A. Offerhaus, Thomas Stervinou, Isabella Mengarelli, Leander Beekman, Lisanne M.W. Wilde, Simona Casini, Tom Streef, Maurice J.B. van den Hoff, Marie-José Goumans, Arie O. Verkerk, Sean J. Jurgens, Arthur A.M. Wilde, Guillaume Lamirault, Nathalie Gaborit, Bastiaan J. Boukens, Carol Ann Remme, Connie R. Bezzina, Fernanda M. Bosada

## Abstract

Brugada syndrome (BrS) is an inherited cardiac arrhythmic disorder caused by conduction slowing primarily affecting the right ventricular (RV) outflow tract (RVOT). A recent genome-wide association study (GWAS) implicated a genomic region in chromosome 11, overlapping the transcription factor *WT1*, in BrS susceptibility. Here, we investigated the role of *WT1* on cardiac conduction using a heterozygous knockout mouse model (*Wt1^+/-^*). Transcriptomic analysis revealed increased *Scn5a* predominantly in *Wt1^+/-^* cardiomyocytes located subepicardially in the RV and RVOT without any changes in electrical properties. To unmask an effect on cardiac conduction, we performed optical mapping in a severely challenged setting offered by *Scn5a* haploinsufficiency, ageing, and exposure to the sodium channel blocker ajmaline and found that diminished *Wt1* improved the observed slowed conduction. Examination of human single-nuclei cardiac datasets indicated a strong negative correlation between *WT1* and *SCN5A* expression. In line with this observation, cardiac samples from patients carrying mutations in *SCN5A* showed increased WT1 protein abundance in histological sections, suggesting that increased *WT1*, and not loss, is associated with BrS pathophysiology. By deleting the mouse orthologue of a BrS-associated noncoding region (RE) harboring a candidate regulatory element, we established that this RE controls expression of *Wt1* specifically in the (sub)epicardium of the RV. Lastly, transient overexpression of *WT1* in hiPSC-derived cardiomyocytes resulted in notably reduced sodium current density. Our study thereby identifies the transcription factor *WT1* as a novel contributor to the pathophysiology of BrS, at least in part, through *SCN5A*.

## Introduction

Brugada Syndrome (BrS) is a heritable cardiac arrhythmia associated with increased risk of sudden cardiac death in young adults^1–3^. The pathophysiological mechanisms underlying BrS remain incompletely understood, hindering development of much needed strategies for risk stratification for life-threatening arrhythmic events and therapy. A long-standing hypothesis revolves around the slowing of conduction of the cardiac electrical impulse, to which the right ventricle (RV) and right ventricular outflow tract (RVOT) are particularly sensitive^4^ likely due to their lower conduction reserve^5^ and propensity for fibrosis^6^. This premise is supported by the fact that rare pathogenic loss-of-function variants in *SCN5A*, encoding the cardiac sodium channel (Na_V_1.5), a critical determinant of cardiac conduction, are found in ∼20% of BrS patients^7,8^ although the vast majority of cases remain genetically elusive^9^. BrS was, until recently, considered a monogenic disease. However, major advances in genetics have uncovered more complex (polygenic) inheritance. Genome-wide association studies (GWAS) have identified 12 genetic loci harboring common variants associated with increased risk for BrS^10,11^. The majority of these disease risk-associated variants are found in noncoding genomic regions, and are presumed to alter the functionality of regulatory elements (REs)^12–14^, which in turn lead to changes in gene expression. The disease-associated loci harbor *SCN5A*, genes that converge at Na_V_1.5 regulation via transcriptional or post transcriptional mechanisms^10,15,16^, as well as transcription factors (TFs) that may play a role in patterning of ion channel expression across the ventricular wall^15^. This supports the notion that diminished Na_V_1.5 function is a central mechanism in BrS pathophysiology. Studies aimed at reliably identifying genes contributing to disease, and exploring how variants affect their expression, are crucial to understand BrS pathophysiology.

GWAS identified a genomic region enriched with noncoding BrS-associated variants in chromosome 11, directly upstream of *WT1* (Wilm’s Tumor 1)^10^. This transcription factor is expressed during development and is known to drive epithelial-to-mesenchymal transition (EMT) to form a population of multipotent epicardial-derived cells^17,18^. Endogenous *Wt1* expression can be found in cardiomyocytes during development^19–22^; while it diminishes postnatally, it can become re-expressed upon injury^23–26^ or in the setting of disease^27–29^. More recently, two studies demonstrated that conditional ablation of *Wt1* in cardiomyocytes during development or postnatally results in morphological abnormalities and *in vivo* electrophysiological changes^22,30^. However, the possible impact of WT1 on ion currents has yet to be explored. Here, we establish a clear association between this TF and BrS, while demonstrating the direction of change in expression that may be present in patients.

## Results

### *Wt1*^+/-^ RV cardiomyocytes show altered ion channel expression without detectable electrophysiological changes

Since GWAS indicated an association with variants directly overlapping *WT1*^10^, we studied the role of this transcription factor using a heterozygous null mouse model (*Wt1^+/-^*). To investigate how reduced *Wt1* affects the transcriptomes of cardiomyocytes, we performed RNA-sequencing on the PCM1+ nuclei from RV/RVOT tissue of aged *Wt1^+/-^*(n=4; 15+ months) mice and age-matched controls (n=4) (Figure 1A). Aged mice were studied in order to resemble the age at which BrS becomes apparent in patients. Among 10722 transcripts detected (>10 averaged normalized reads), 466 were upregulated and 405 downregulated in *Wt1^+/-^* cardiomyocyte nuclei (Figure 1B). Notably, *Wt1* transcripts were almost undetectable in our cardiomyocyte-enriched samples (Supplemental Table 1). Gene ontology term analysis using PANTHER^31^ identified biological processes including development (*Rbm20*, *Mef2d*, *Tcf7l2* and *Sox6*), cardiac muscle contraction (*Cacna1c*, *Ryr2*, *Slc9a1* and *Myh7b*), and cell motility (*Actg1*, *Ctnna1*, *Cdh13* and *Sema6d*) associated with upregulated genes in *Wt1^+/-^*. Cellular respiration (*Mtch2*, *Cox7b*, *Mdh1* and *Dld*), muscle tissue development (*Tgfb2*, *Myoz2*, *Hand2* and *Gata6*), and intracellular transport (*Dnm1l*, *Mtx2* and *Calm1/2/3*) were associated with the downregulated gene set (Figure 1C). Strikingly, we found increased *Scn5a* (L2FC 0.2333, p-adj=0.0124) and various deregulated genes involved in cellular electrophysiology in *Wt1^+/-^* cardiomyocytes (Figure 1B, D).

**Figure 1.**
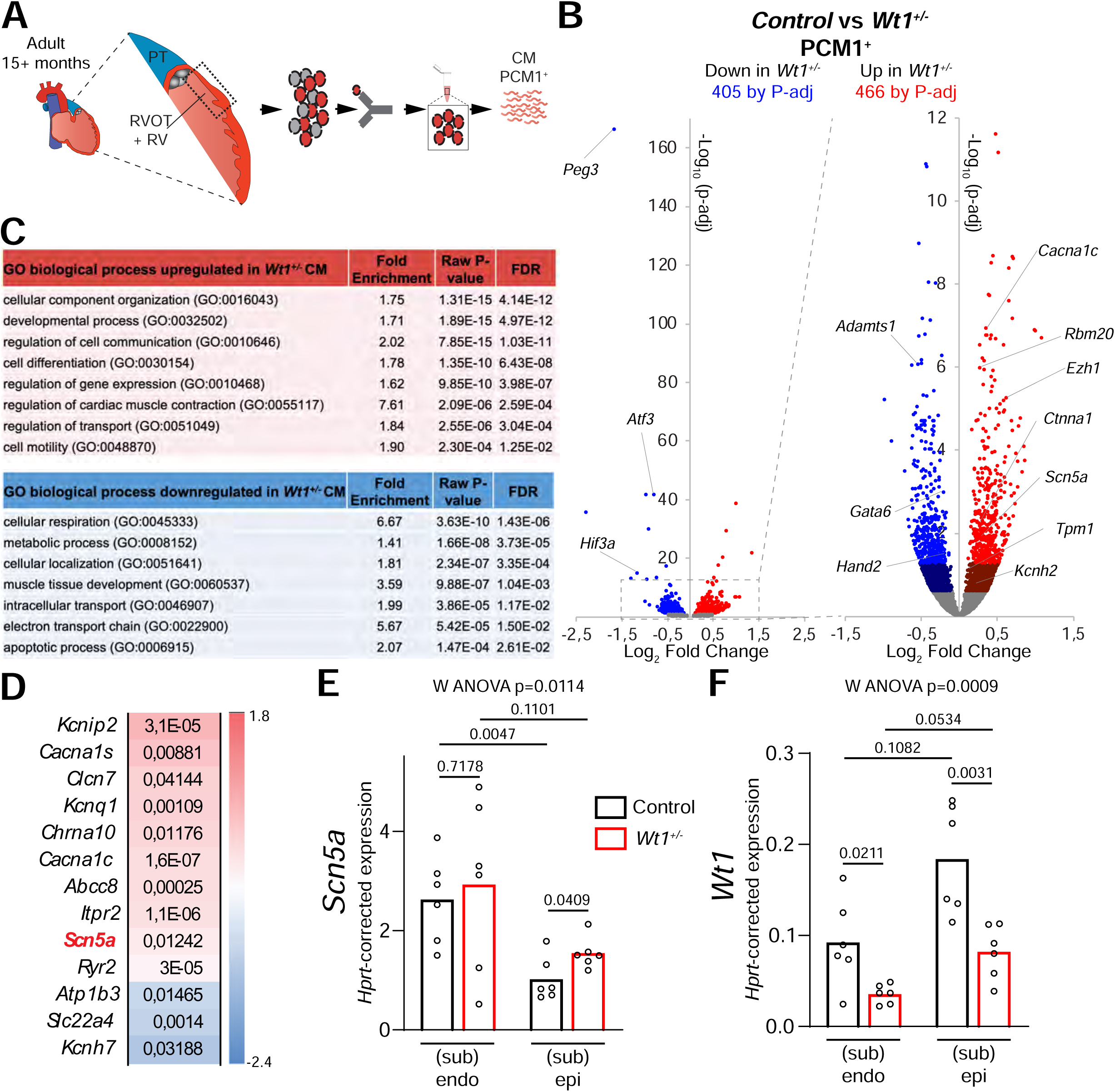
Diminished *Wt1* leads to increased *Scn5a*. A) Schematic depicts approach for isolating cardiomyocyte (CM) nuclei of the RV and RVOT. B) Volcano plot shows differentially expressed transcripts between control (n=4) and *Wt1^+/-^*(n=4) cardiomyocytes. Dark red and light red dots indicate significantly upregulated genes by raw p-value and p-adjusted for multiple testing, respectively. Correspondingly, dark blue and light blue dots indicate downregulated genes. C) Gene ontology (GO) analysis of up-and downregulated genes in *Wt1^+/-^* cardiomyocytes. P-values were adjusted for multiple testing using the false discovery rate (FDR) method of Benjamini-Hochberg. D) Heatmap depicting deregulated genes and their adjusted p-values (all p<0.05) involved in electrical conduction. Scale denotes fold change. E, F) *Hprt*-corrected expression of *Scn5a* (E) and *Wt1* (F) from adult control (n=6) and *Wt1^+/-^* (n=6) ventricular (sub)endocardium and (sub)epicardium determined by RT-qPCR. Significant p-values were determined with Welch’s ANOVA followed by unpaired *t*-tests with Welch’s correction.

Given the well-established ventricular transmural ion channel expression gradient for normal conduction^15,32^, we tested the impact of diminished *Wt1* in the (sub)endocardial and (sub)epicardial sides of the ventricular wall on *Scn5a* by RT-qPCR. As expected, we observed that in control hearts, *Scn5a* expression was higher in the (sub)endocardial side of controls (Figure 1E), while *Wt1* showed a clear tendency to higher expression in the (sub)epicardial side (Figure 1F). We also confirmed that *Wt1* was significantly downregulated in *Wt1^+/-^*ventricles in both transmural compartments (Figure 1F). Remarkably, we detected increased *Scn5a* specifically in the (sub)epicardial side of *Wt1^+/-^* ventricles when compared to controls (Figure 1E). This indicates that the increase in *Scn5a* expression detected in cardiomyocytes of the RV/RVOT of *Wt1^+/-^* mice is driven by a mechanism primarily present in the (sub)epicardial side of the ventricle, thus suggesting that epicardial *Wt1* inhibits *Scn5a*.

ECG analyses of adult (3-4 months) mice did not show any differences in PR-, QRS-and QT duration in *Wt1^+/-^* mice compared to age-matched controls (Supplemental Figure 1A), indicating no overt differences in cardiac electrophysiological properties. RV and RVOT (sub)epicardial conduction was measured in more detail in Langendorff-perfused hearts of adult mice using optical mapping (Supplemental Figure 1B, C). Overall, no differences in conduction velocity (CV) were observed between *Wt1^+/-^* and control mice in either the RV or the RVOT. To study the effect of *Wt1* haploinsufficiency on cellular electrical properties we measured action potentials (AP) and sodium current (I_Na_) in transmurally unselected cardiomyocytes isolated from the RV of adult control and *Wt1^+/-^* mice. AP upstroke velocity (V_max_) was similar in cardiomyocytes of both groups (Supplemental Figure 2A, B, Supplemental Table 1). In line with this, no differences were observed in I_Na_ density or voltage dependence of (in)activation (Supplemental Figure 2C-E, Supplemental Table 2). Resting membrane potential (RMP), AP amplitude (APA) and AP repolarization parameters APD_20_, APD_50_ and APD_90_ were also not significantly different between RV cardiomyocytes from adult control and *Wt1^+/-^* mice (Supplemental Figure 2A, B, Supplemental Table 2). Together, our data indicate that diminished *Wt1* expression results in gene expression changes which are not sufficient to lead to an electrophysiological phenotype.

### Diminished *Wt1* improves conduction in a severely challenged setting

Next, we evaluated the possibility that electrophysiological changes driven by diminished *Wt1* may only become apparent in the context of reduced conduction. To investigate this, we crossed *Wt1^+/-^* mice with *Scn5a* haploinsufficient mice (*Scn5a^+/-^*) to generate double haploinsufficient mice (*Scn5a^+/-^*;*Wt1^+/-^*) and examined whether aging provides a sufficiently sensitized setting. We therefore aged control, *Scn5a^+/-^* and *Scn5a^+/-^*;*Wt1^+/-^*mice to ∼15 months, equivalent to the approximate age at which BrS is diagnosed on average in humans (40-50 years)^2,33^. *In vivo* ECGs of aged mice (Supplemental Figure 3A) revealed that *Scn5a* haploinsufficiency led to prolonged PR interval (Supplemental Figure 3B), while QRS duration was unchanged, regardless of *Wt1* loss (Supplemental Figure 3C). Aging, independent of genotype, did not have an effect on PR duration but did result in QRS interval prolongation in both *Scn5a^+/-^*and *Scn5a^+/-^*;*Wt1^+/-^*, but not in control mice (Supplemental Figure 3). Indeed, an interaction effect was borderline significant for QRS duration (p=0.0656), suggesting that aging may unmask a more pronounced ventricular conduction phenotype in mice with only one copy of *Scn5a.* We next investigated this in more detail by measuring epicardial CV in the RV and RVOT in aged mice. We found slower CV in both longitudinal and transverse direction in *Scn5a^+/-^* and *Scn5a^+/-^*;*Wt1^+/-^*RVs than in controls (Figure 2A). In the RVOT we observed slower conduction in *Scn5a^+/-^* compared to controls, but this difference was not detected between *Scn5a^+/-^*;*Wt1^+/-^*and controls (Figure 2B). We then further challenged our experimental model (aged *Scn5a^+/-^*and *Scn5a^+/-^*;*Wt1^+/-^* animals) by suppressing I_Na_ availability through acute administration of the I_Na_ blocker ajmaline. Since the relationship of CV and I_Na_ is not linear^34^, CV differences are more easily to detect by the reduced excitability. Optical mapping analysis in the presence of ajmaline showed a significantly higher CV in the transverse direction of the RV (Figure 2C), and in the RVOT (Figure 2D) of *Scn5a^+/-^*;*Wt1^+/-^* compared to *Scn5a^+/-^*mice. Taken together, our data show that ageing contributes to ventricular conduction delay, and that decreased *Wt1* expression improves conduction in *Scn5a* haploinsufficient mice in the setting of severely reduced conduction reserve.

**Figure 2.**
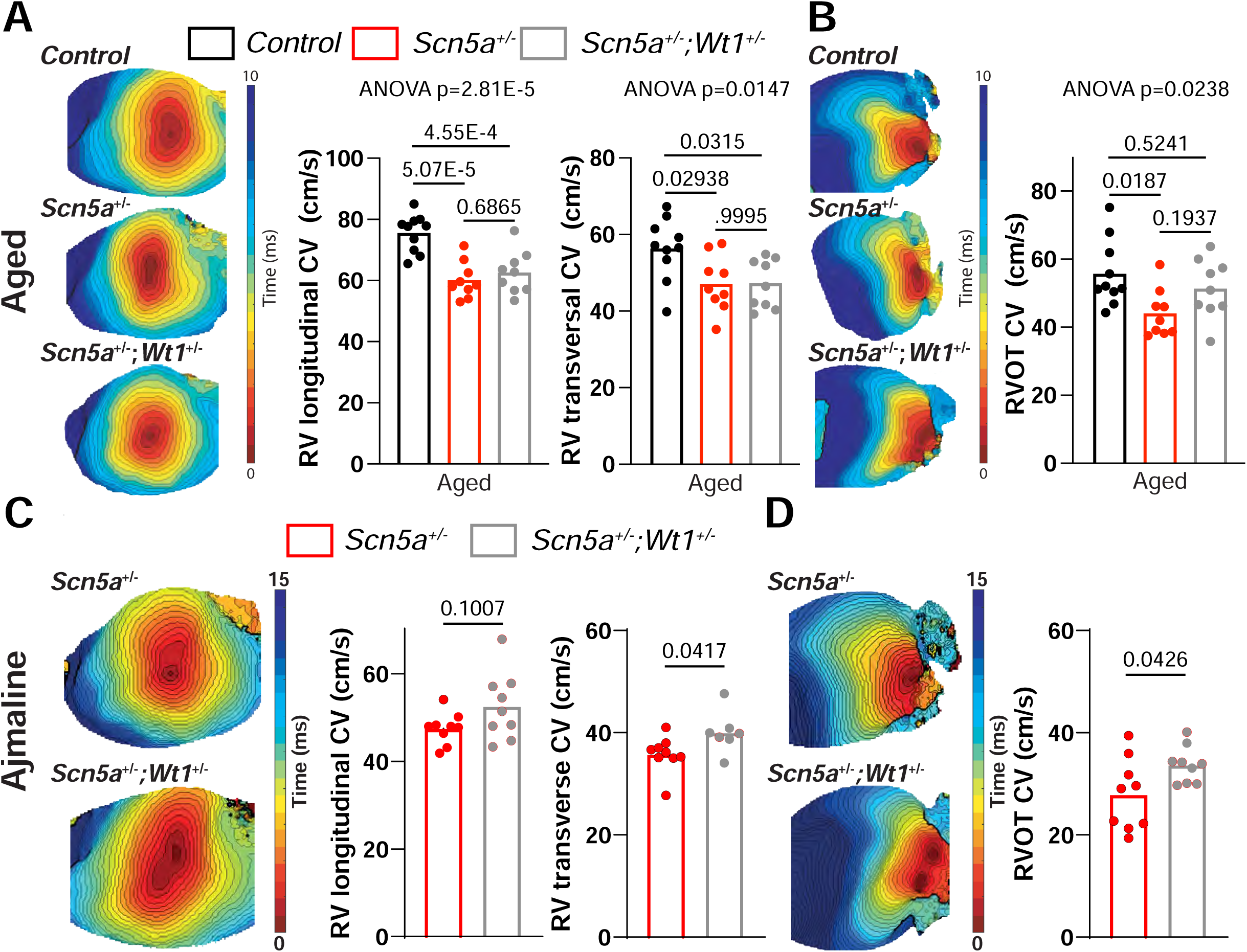
Reduced *Wt1* expression improves conduction in the setting of severely reduced conduction in aged *Scn5a^+/-^* mouse hearts. A) Typical examples of RV longitudinal and transversal-and B) RVOT conduction with respective average conduction velocity (CV) of aged control (n= 10), *Scn5a^+/-^*(n= 9) and *Scn5a^+/-^*;*Wt1^+/-^* (n= 9). RV, right ventricle; RVOT, right ventricular outflow tract. C) Typical examples and average CV of RV longitudinal and transversal direction, and D) RVOT in aged *Scn5a^+/-^* and *Scn5a^+/-^*;*Wt1^+/-^*mice after acute administration of ajmaline (n=7). Statistical significance was determined with ANOVA followed by pairwise Tukey’s multiple comparison tests for A and B, and unpaired *t*-tests for C and D.

### Inverse correlation between *WT1* and *SCN5A* expression in human transcriptomic datasets

Because diminished *Wt1* leads to increased *Scn5a* in cardiomyocytes, as well as a rescue effect on conduction slowing in *Scn5a* deficient mice, we explored whether a correlation between *WT1* and *SCN5A* expression is present in human cardiac samples. Using GTEx data, we defined two groups of samples, classified as “Low” or “High” cardiomyocyte-enriched, based on 10 cardiomyocyte-specific genes^35^, which are significantly correlated with each other (Supplemental Figure 4A-C). In “High” cardiomyocyte-enriched samples, we observed a significant negative correlation between *SCN5A* and *WT1*, whereas no correlation was present in the “Low”, non-cardiomyocyte-enriched group (Supplemental Figure 4D). This correlation was further supported by single-cell RNA-seq (snRNAseq) data of human induced pluripotent stem cell-derived cardiomyocytes (hiPSC-CMs) (Figure 3A, B). We then assessed the relationship between *WT1* and *SCN5A* expression in all atrial and ventricular cardiomyocytes (Figure 3C), and in atrial and ventricular cardiomyocytes that express either, or both of these transcripts (Figure 3D). We found a strong negative correlation, which indicates that cells with higher *WT1* expression have tapered *SCN5A* and vice-versa and this may thereby impact on the transmural expression gradient observed in mice (Figure 1E, F). We confirmed this result in snRNAseq datasets of non-diseased human hearts^28^. Pseudo-bulk regression analysis across 4 ventricular regions revealed a strong negative correlation between *WT1* and *SCN5A* (Supplemental Figure 4E, F, Supplemental Table 3).

**Figure 3.**
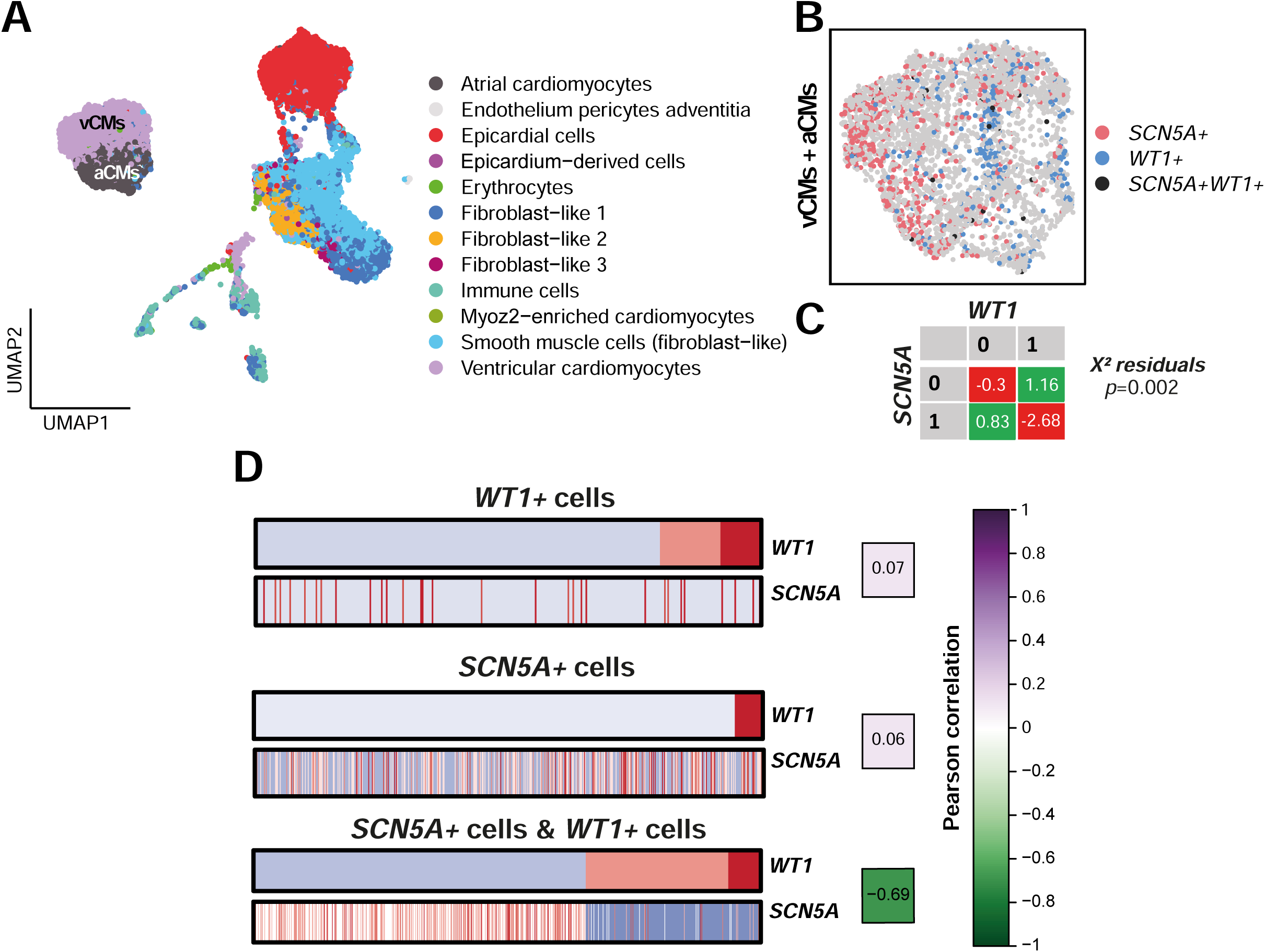
A negative correlation between *WT1* and *SCN5A* expression. A) UMAP of single-cell RNA-seq data from control hiPSC-CMs, annotated using a 7 post-conception week (PCW) human embryonic heart reference dataset, published by^76^. Cardiomyocytes (CMs) are displayed on the UMAP as ventricular (vCMs) or atrial (aCMs). B) UMAP of vCMs and aCMs. Cells expressing *SCN5A* only, *WT1* only, and both *SCN5A* and *WT1* are displayed in pink, blue, and black, respectively. C) Contingency table showing the Chi-square test residuals for cells expressing (1) or not expressing (0) *SCN5A* and *WT1*. Negative and positive residual values indicate that the corresponding cell proportions are under-or over-represented compared to the expected distribution, respectively. D) Pearson correlation analysis between *SCN5A* and *WT1* expression in all *WT1*+ cells (top), all *SCN5A*+ cells (middle), and cells expressing at least one of the two genes (*bottom*). Cells are ordered by increasing *WT1* expression: blue, indicates absence of expression, red, indicates highest level of expression and pink, indicates intermediate level of expression.

Several of the conduction-related genes altered in cardiomyocytes of *Wt1^+/-^* mice, including *Scn5a* (Figure 1D), are similarly correlated with *WT1* expression in human left ventricular samples (GTEx). *WT1* is negatively correlated with *SCN5A, RYR2, KCNQ1, CACNA1C* and *KCNIP2,* whereas it is positively correlated with *ATP1B3* (Supplemental Figure 5). This evidence indicates that *Wt1* haploinsufficiency in mice mirrors the transcriptional state observed in human heart tissue, thereby strengthening the translational relevance of mouse models.

### *WT1* is increased in pathologic conditions including BrS

In normal hearts, *WT1* expression diminishes after birth, but it can become re-expressed upon injury^23–26^. Recent snRNAseq studies have shown increased *WT1* in ventricular samples of patients with congenital cardiac defects and hypertrophic cardiomyopathy (HCM) when compared to non-diseased hearts^27–29^. Moreover, patients with arrhythmogenic cardiomyopathy (ACM), a disease exhibiting overlapping pathology with BrS, showed increased *WT1* when compared to healthy controls^36^. We then considered whether elevated *WT1* expression occurs in the context of BrS. To test this, we obtained histological ventricular sections from three *SCN5A* mutation-positive patients. Immunohistological staining for WT1 revealed distinct nuclear signal in 2 out of 3 of these hearts, while non-diseased hearts, neither young (23 years) or old (79 years), showed barely any detectable staining (Supplemental Figure 6). These data indicate that increased *WT1*, and not its loss, is consistent with the BrS phenotype.

Next, we considered whether *SCN5A* loss-of-function itself leads to induction of *WT1* at any point in time. To address this, we performed lineage tracing experiments using *Wt1^Cre^*^18^*;R26R-mTmG*^37^, in which the Cre recombinase drives membrane GFP only in the *Wt1* lineage, and studied this in the context of *Scn5a* haploinsufficiency. We quantified GFP+ cardiomyocytes of the RV, left ventricle (LV), and interventricular septum (IVS) of aged *Scn5a^+/-^* and control mice using TnT staining and flow cytometry. We found significantly more GFP+ cardiomyocytes in the IVS and LV of *Scn5a^+/-^* mice when compared to controls (Figure 4A). We next analyzed GFP+/cTnT+ cells on histological sections and found increased double positive cells in mice with reduced *Scn5a* expression (Figure 4B, C). These results indicate that life-long reduced *Scn5a*, as it occurs in patients with loss-of-function *SCN5A* mutations, leads to induction of *Wt1* in cardiomyocytes at any point through life, which further solidifies the biological relevance of the inverse relationship between *Wt1* and *Scn5a*.

**Figure 4.**
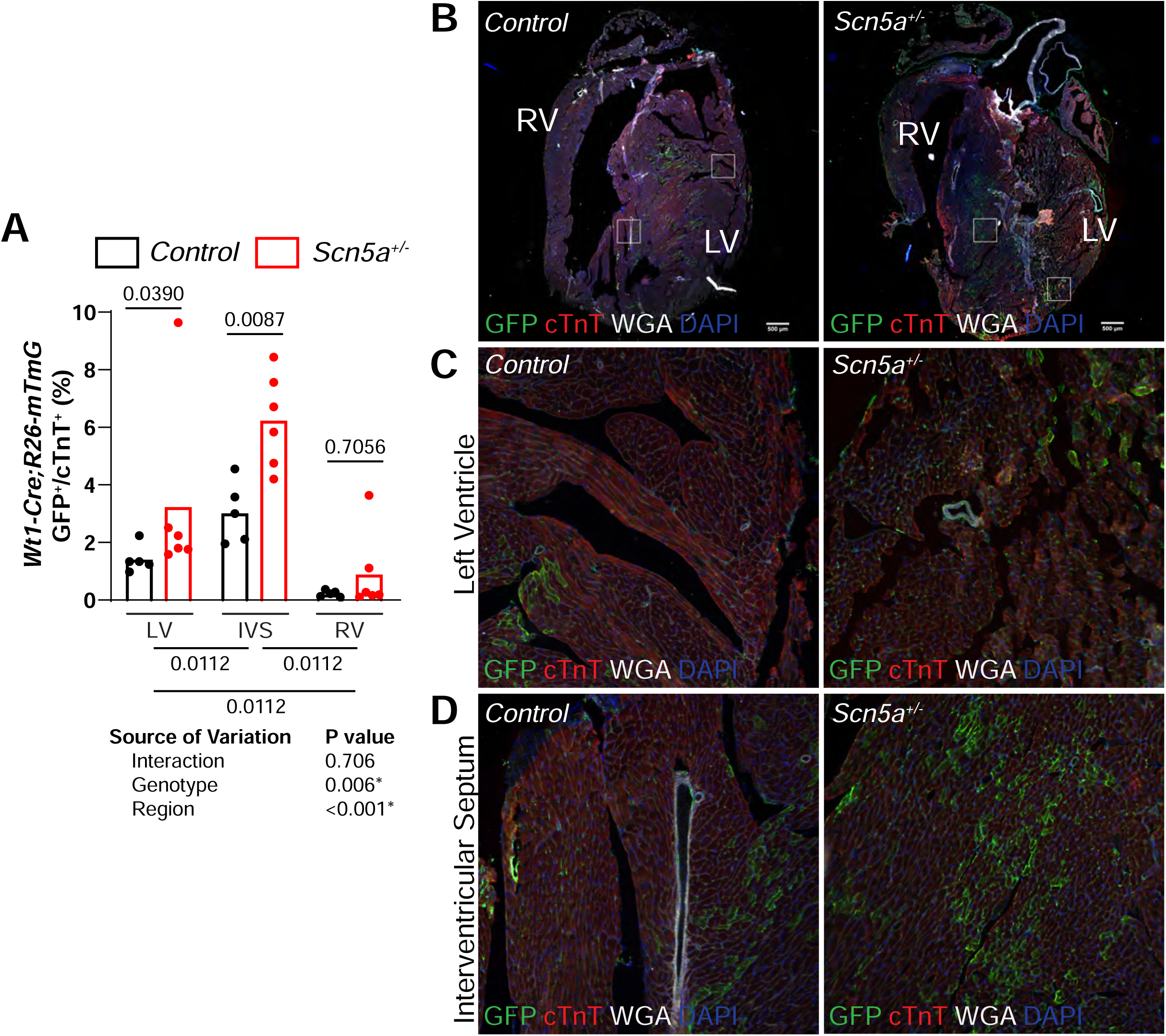
Increased *Wt1*-lineage cells in *Scn5a* haploinsufficient hearts. A) Percent GFP+/cTnT+ cells in LV, IVS, and RV in control (n= 5) and *Scn5a^+/-^* (n=6) mice as determined by FACS. Significance determined by a general linear mixed model and post-hoc tests. B-D) Representative images of histological sections of control and *Scn5a^+/-^* hearts. LV, left ventricle; RV, right ventricle; IVS, interventricular septum.

### The BrS-associated noncoding variant region controls *Wt1* expression in the RV

Given that diminished *Wt1* expression may confer a protective effect, and patients with *SCN5A* loss-of-function mutations display elevated WT1, we proceeded to investigate how the BrS-associated variant region (BAR) in chr11p13 impacts on *WT1.* We first identified candidate regulatory elements within this region. Human epigenetic datasets, including ventricular-like hiPS cardiomyocyte ATACseq (a measure of open chromatin regions)^38^, CTCF ChIPseq^39^, and the EMERGE enhancer prediction track^40,41^ reveal moderate epigenetic signatures overlapping the BrS-associated variant region (Figure 5A). In the mouse orthologous region, ATACseq of adult ventricular and neonatal sinus node tissue^42,43^ shows open chromatin in the region upstream of *Wt1*, with modest H3K27ac signatures^44^ (Figure 5B). We therefore selected a 4903 bp region overlapping these epigenetic signatures for deletion using CRISPR/Cas9 (Figure 5C). Homozygous deletion mice (*Wt1^BAR/BAR^*) were born at the expected Mendelian ratios and showed no overt morphological defects. Using RTqPCR, we determined the expression level of *Wt1* in the (sub)epicardial and (sub)endocardial regions of RVs and LVs of 6-week-old mice. *Wt1^BAR/BAR^* mice showed a ∼45% increase in *Wt1* transcripts in the (sub)epicardial side of the RV, but not in any other cardiac region assayed (Figure 5D). This observation indicates that the BAR contains one or more regulatory elements that selectively represses expression of *Wt1* in the (sub)epicardium of the RV, thus further supporting the notion that increased *WT1* contributes to BrS pathophysiology.

**Figure 5.**
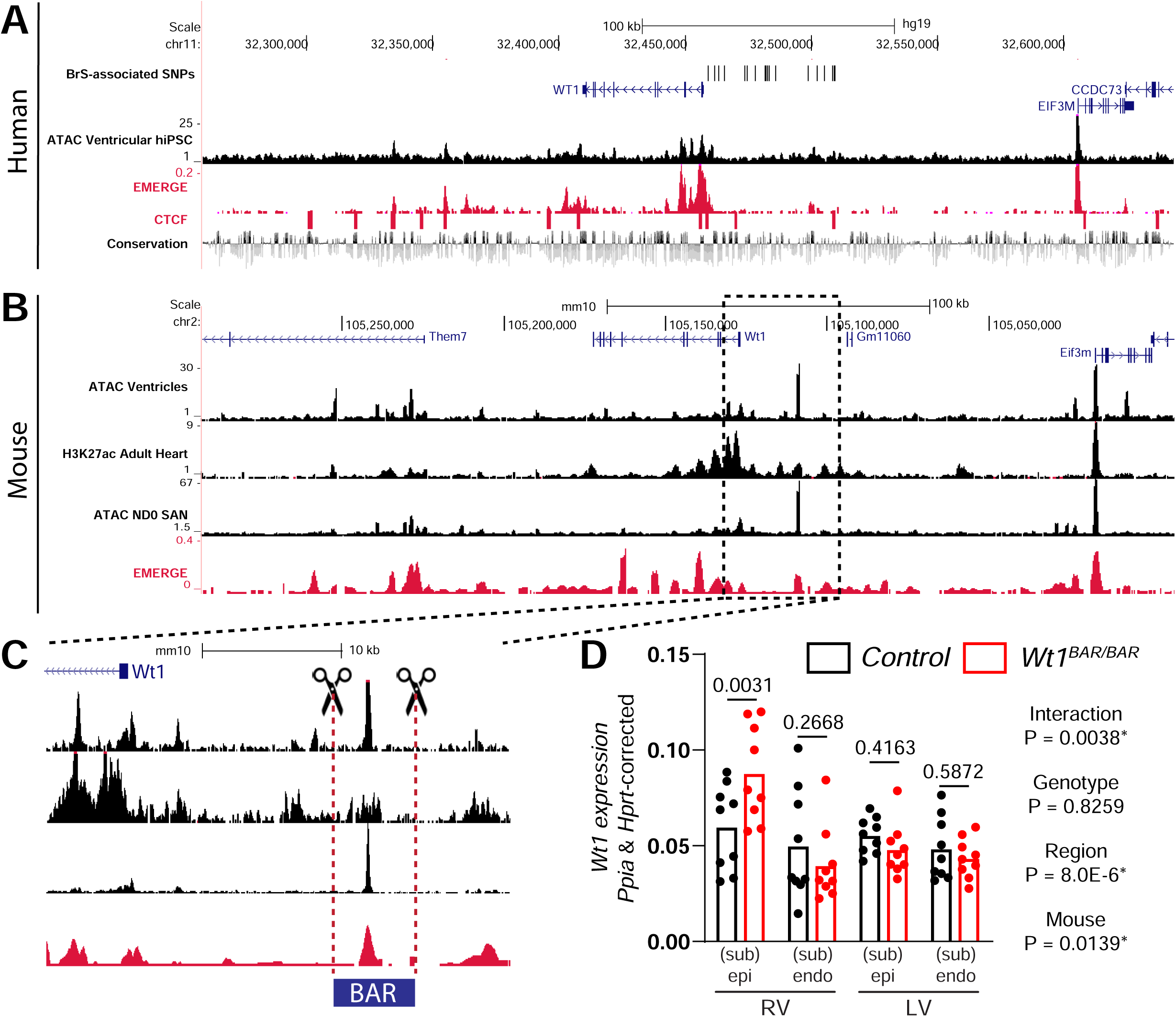
Deletion of BrS-associated region in mouse orthologue leads to increased *Wt1* selectively in (sub) epicardium of RV. A) UCSC browser view of the human BrS-associated locus in chr11 including ATACseq from hiPS ventricular cardiomyocytes (black), EMERGE (red), CTCF ChIPseq (red bars), and conservation track. B) Mouse orthologue of the human region shown in panel B including ATACseq of ventricular cardiomyocytes and Hcn4+ sinoatrial node (SAN) cells, H3K27ac ChIPseq from whole adult hearts, and EMERGE. C) Zoom-in of panel B depicting the deleted region (BAR) in blue. D) Reference gene-corrected expression of *Wt1* from isolated (sub) epicardial and (sub)endocardial sides of the RV and LV determined by RTqPCR of *Wt1^BAR/BAR^* (n=9) and control littermates (n=9). Statistical significance was determined with 2-way ANOVA followed by pairwise comparisons with Tukey’s multiple comparison tests. P values are shown.

### Overexpression of WT1 in hiPSC-derived cardiomyocytes leads to reduced sodium current

To determine the functional consequences of *WT1* overexpression in single cardiomyocytes, we generated hiPSC-CMs by using an adapted protocol based on small molecule-mediated modulation of the cWnt-pathway^45^. hiPSC-CMs were transfected using Lipofectamine 3000 with plasmids containing either WT1-P2A-GFP or GFP only (which served as control) and measured peak sodium (I_Na_) in GFP-positive (GFP^+^) hiPSC-CMs. Peak I_Na_ density was significantly lower in hiPSC-CMs transfected with WT1 (Figure 6A, B). For example, I_Na_ density at –20 mV was-87±10 and-54±7 pA/pF (*p*<0.05; t-test) in GFP^+^ (n=16) and WT1 (n=12) overexpressing hiPSC-CMs. Neither the voltage dependency of (in)activation nor the time constants of current inactivation, determined at a test potential of-20 mV, were significantly different in hiPSC-cardiomyocytes with WT1 overexpression (Figure 6C, Supplemental Table 4), indicating that WT1 overexpression only affects I_Na_ density and not the gating properties. While a previous study reported that loss of *Wt1* in embryonic cardiomyocytes led to dysregulation of a potassium channel subunit which is an important regulator of the transient outward (I_to1_) current^22^, we did not detect any differences in I_to1_ density (Figure 6D, E) or I_to1_ gating properties (Figure 6F, Supplemental Table 4). AP measurements performed using dynamic clamp^46^ demonstrated a significant decrease in V_max_ in hiPSC-CMs overexpressing WT1 compared to controls (Figure 6G, H), in line with the observed reduced I_Na_. The RMP, APA, and AP repolarization parameters remained unchanged upon WT1 overexpression (Figure 6H, Supplemental Table 4). Taken together, these observations align with the effect of *Wt1* reduction on ventricular conduction demonstrated herein, and suggest a modulatory contribution predominantly on depolarization.

**Figure 6.**
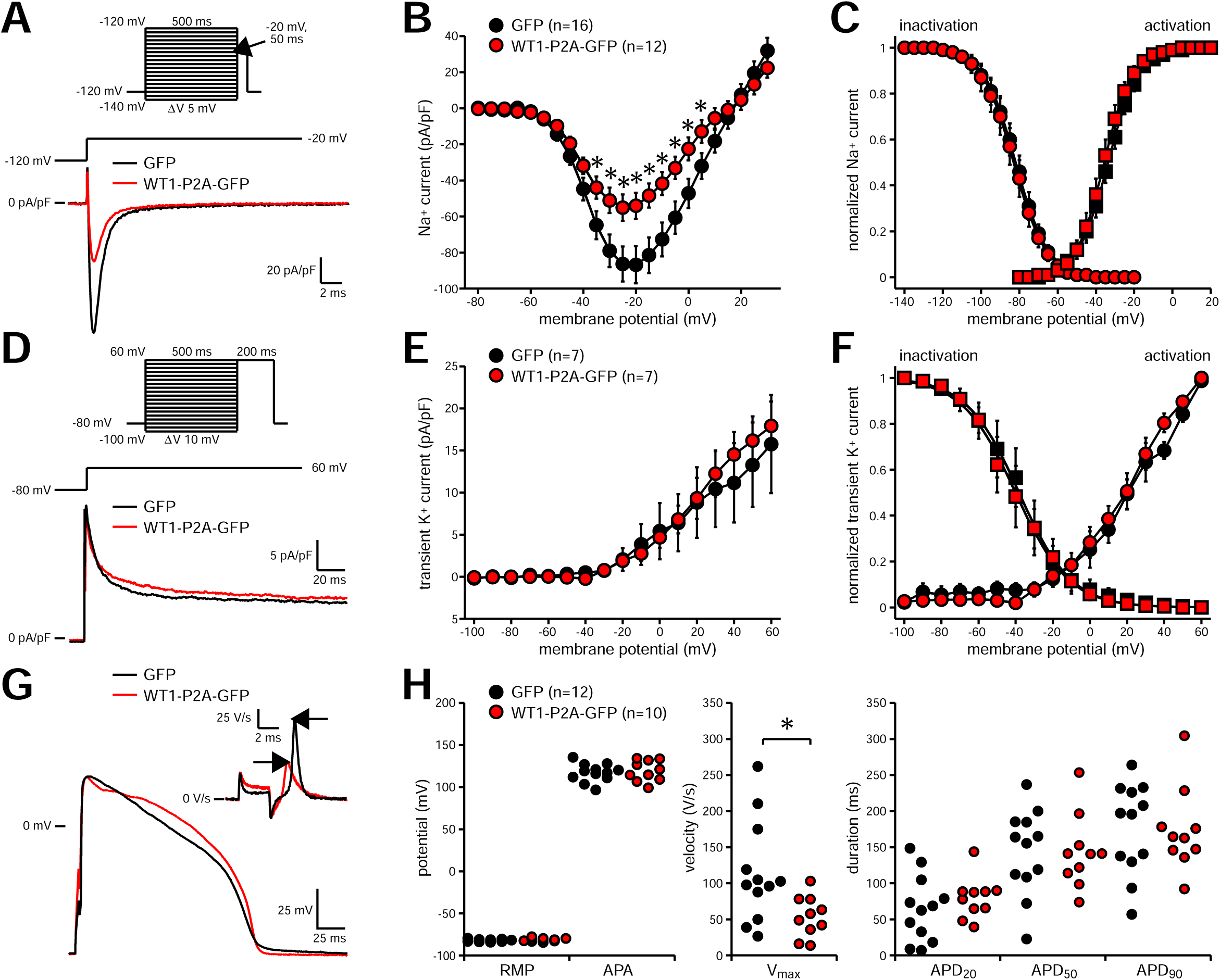
WT1 overexpression in hiPSC-cardiomyocytes reduces sodium current (I_Na_) and action potential (AP) upstroke velocity. A) Voltage clamp protocol and representative traces of I_Na_ activated during depolarizing steps from-120 to-20 mV. B) Average current-voltage (I-V) relationships of I_Na_. C) Average voltage dependence of (in)activation curves. Solid lines represents the fitted Boltzmann function (I/I_max_=A/{1.0+exp[(V_1/2_ −V)/k]}). D) Voltage clamp protocol and representative traces of the transient outward potassium current (I_to1_) activated during depolarizing steps from-80 to 60 mV. E) Average I-V relationships of I_to1_. F) Average voltage dependence of (in)activation curves. Solid lines represent the fitted Boltzmann function for inactivation. G) Typical APs measured at 1 Hz. The inset shows the <u>AP upstroke velocity</u> (V_max_). **H.** Average AP parameters. RMP, resting membrane potential; APA, AP amplitude; APD_20_ APD_50_, and APD_90_, AP duration at 20, 50 and 90% of repolarization, respectively; * Indicates P<0.05 by Two-Way ANOVA or unpaired t-test.

## Discussion

Here, we explored the role of *WT1*, a gene at a locus implicated in BrS pathophysiology by GWAS. Our study reveals that diminished *Wt1* expression in mice leads to altered ion channel and cardiac contraction gene expression, including a modest increase in *Scn5a* specifically in subepicardial cardiomyocytes of the right ventricular wall. We further provide evidence supporting an inverse relationship between *WT1* and *SCN5A* expression in humans and identified increased *WT1* in pathological conditions, including in *SCN5A* mutation positive BrS patients. The interaction between these two genes was further explored in a challenged setting provided by *Scn5a* haploinsufficiency. While no differences in electrophysiological properties either *in vivo*, *ex vivo*, or in single cardiomyocytes were detected under baseline conditions, diminished *Wt1* did improve conduction under conditions of suppressed conduction reserve in the setting of advanced age and acute ajmaline exposure. By deleting the BrS-associated noncoding region orthologue, we show that *Wt1* is a target of the REs likely affected by the disease-associated common variants. Lastly, we demonstrate that overexpression of WT1 in hiPSC-derived cardiomyocytes reduces I_Na_ density, thereby establishing *WT1* as a new player in the pathology of BrS.

A recent GWAS proposed *WT1* as the most likely candidate gene implicated in BrS in chromosome 11, as indicated by the association signal in this locus^10^. In our study, we demonstrate that lifelong reduction of *Wt1* leads to an upregulation of *Scn5a* predominantly on the subepicardial side of the cardiac ventricles. This finding points to a potential mechanism by which *WT1* contributes to BrS through regulation of *SCN5A*, while also suggesting that increased-, rather than decreased WT1 expression, is associated with the BrS pathological state. Moreover, this highlights *WT1* as a potential key regulator in maintaining the transmural ion channel gradient, which is essential for proper cardiac function. Correspondingly, we demonstrate that patients with *SCN5A* mutations exhibit elevated WT1 protein levels in cardiac tissue, and that transient overexpression of WT1 in hiPSC-derived cardiomyocytes results in a significant reduction in sodium current. This is consistent with existing knowledge regarding the loss of Na_V_1.5 function in the pathophysiology of BrS.

From an *in vivo* physiological perspective, discerning the impact of reduced *Wt1* is challenging. In order to observe an electrophysiological phenotype, it was necessary to challenge WT1 haploinsufficient mice by aging and reducing sodium channel expression and activity through acute exposure to ajmaline. This could be due to the complexity associated with accurately measuring increased Na_V_1.5 function in a laboratory setting. *Scn5a^+/-^* mice showed increased QRS duration only with ageing, and not at young adult age as was shown before^47^. A possible explanation for the mitigated phenotype is differences in used mouse strain, affecting electrophysiological parameters^48,49^. In this context, a rescue effect provided by *Wt1* reduction (and thereby increased *Scn5a*) can only be uncovered by aggravating the phenotype even more. Together, these observations provide strong evidence that *WT1* impacts on SCN5A expression, and thereby also BrS pathophysiology.

The BrS-associated common variants in chromosome 11 are located in the noncoding region upstream of *WT1*, and are therefore thought to impact on gene expression via REs^12–14^. RE activity is largely limited to the topologically-associating domain they share with their target genes^50–52^. We presume that BrS-risk variants disrupt the function of REs that control *WT1* expression. Consequently, by deleting the mouse orthologue of the candidate RE, we aim to model the genetic mechanism(s) at play in BrS patients. Indeed, we demonstrated that loss of the BrS-associated variant-rich RE led to increased *Wt1* specifically in the (sub)epicardial side of the RV *in vivo*. This finding further supports the notion that common variants associated with increased BrS risk at this locus likely result in *WT1* upregulation, which results in loss of Na_V_1.5 function, thereby contributing to increased BrS-risk. However, further research is required to elucidate the transcriptional regulatory mechanisms driving the function of this and other potential REs within the locus.

We speculate that decreased or increased *WT1* impacts on *SCN5A* expression in the opposite direction via a paracrine or cell-autonomous mechanism. Although WT1 is known to function as either a transcriptional activator or repressor depending on its binding partners (reviewed in^53^), not all evidence supports a cell-autonomous mechanism. We show that *Wt1* expression was below detection in the cardiomyocyte-enriched fraction, yet *Scn5a* was upregulated in *Wt1^+/-^*cardiomyocytes. This result may indicate that deregulation of *WT1* in non-cardiomyocytes, via a paracrine process, affects gene expression in cardiomyocytes. The specific cell-type in question has yet to be identified. Alternatively, *WT1* may be expressed at low levels, in a small proportion, or only transiently in cardiomyocytes, making it difficult to detect any decrease in expression, and thereby introducing further challenges for the interpretation of data. Our study, along with others^27–29,36^, indicates that elevated *WT1* expression is linked to pathological states. Therefore, only in the context of increased WT1 we may be able to uncover a cell-autonomous transcriptional effect on *SCN5A*.

It is also a possibility that WT1 affects SCN5A via a non-transcriptional mechanism. Recent studies have revealed that WT1 can also control gene expression by modulating the epigenetic landscape via interacting with TET family proteins^54,55^. WT1 is also known to modulate topological switch mechanisms involving global changes in chromatin access to provide context for specific gene activation or repression. For example, it acts as a transcriptional activator of *Wnt4* in the developing kidney mesenchyme undergoing mesenchymal-to-epithelial transformation, and a repressor of *Wnt4* in epicardial cells poised for EMT^56^. WT1 may also function post-transcriptionally according to a study which showed that WT1 localizes and interacts with splicing factors in kidney cells^57^. Furthermore, WT1 has been shown to regulate mitotic checkpoint via direct interaction with the spindle assembly checkpoint protein MADS2^58^. Thus, additional research is needed to rule out the possibility of a non-transcriptional contributing mechanism.

Conversely, decreased *SCN5A/Scn5a* expression also results in induction of *WT1/Wt1*, as demonstrated by *Wt1*-lineage tracing on *Scn5a^+/-^* mice and histological staining of patient hearts with an *SCN5A* mutation. This suggests a tightly regulated interplay and the dynamic balance influences cellular processes including cardiac conduction. Once either of these genes shifts towards a pathologic state (i.e. increased *WT1* or decreased *SCN5A*), it may trigger a potential maladaptive cyclic mechanism which progressively exacerbates organ dysfunction.

In conclusion, our data establish a role for the transcription factor *WT1* in the pathophysiology of BrS, at least in part, through modulating expression and function of the cardiac sodium channel *SCN5A*. Additionally, we present an explanation for the association of the noncoding region in chromosome 11 with BrS in patients. Beyond rare coding loss-of-function variants in *SCN5A*, we expect that the aggregate effect of common variants at one or more disease-associated loci impact on Na_V_1.5 function and thereby BrS susceptibility.

## Methods

### Ethics statement

Housing, husbandry, all animal care and experimental protocols were in accordance with guidelines from the Directive 2010/63/EU of the European Parliament, and Dutch government. Protocols were approved by the Animal Experimental Committee of the Amsterdam University Medical Centers. Animal group sizes were determined based on previous experience.

### Mice

*Scn5a^+/-^* and *Wt1^Cre^* mice were generated as described previously^18,47^, respectively. 129S4/SvJae *Wt1^GFPCre/+^* and *R26-mTmG* mice were obtained from Jackson Laboratories (stock numbers 010911, 007676, respectively). Given that the GFP-Cre insertion in the *Wt1* allele replaces exon 1 and thereby produces a null allele^59^, the mice are herein referred to as *Wt1^+/-^*. *Wt1^+/-^* mice were crossed with *Scn5a^+/-^* to generate *Scn5a^+/-^*;*Wt1^+/-^*. *Wt1^Cre^* mice were crossed first to *R26-mTmG* followed by *Scn5a^+/-^* for lineage tracing experiments.

*Wt1^BAR/BAR^* mice were generated using CRISPR/Cas9. Guide RNA (sgRNA) target sites were designed with PrimerQuest™ program (IDT, Coralville, Iowa, USA. Accessed 12 December, 2018. https://www.idtdna.com/SciTools). The Alt-R® CRISPR-Cas9 crRNA, Alt-R® CRISPR-Cas9 tracrRNA (IDT #1072532) and Alt-R® S.p. Cas9 Nuclease V3 (IDT #1081058) were ordered at IDT. One-cell FVB/NRj zygotes were microinjected via the “Mouse zygote microinjection Alt-R™ CRISPR-Cas9 System ribonucleoprotein delivery protocol” with 20 ng/µl Cas9 Nuclease and 10 ng/µl guide RNA to generate mouse founders. The guide sequences were as follows: 1) GGATTTAGGTCAAGATTCAC, 2) CACCACCATGGGATTCCTAA. Deletions were validated by PCR using the following primers: Forward 1) CTCTGGCTGCTGAAGGGATT, Reverse 1) CTAGCTCTTACTGGCCACCG, Reverse 2) GAGAACAGAGCCCAGGAAGG and Sanger sequencing. The deleted region spans chr2:105107107-105112009. To obtain stable lines, the founders were backcrossed with wild-type FVB/NJ mice and maintained on FVB/NJ background ordered from the Jackson laboratory (#100800). For tissue harvest, animals were euthanized by 20% CO_2_ inhalation followed by cervical dislocation.

### Isolation of cardiomyocyte nuclei

Nuclei isolation was performed as previously described^60^ with the following changes: snap frozen adult left and right atria from adult male and female mice were trimmed and homogenized in lysis buffer containing RNase inhibitor using an Ultra-Turrax homogenizer. Samples were further homogenized with a loose pestle douncer (10 strokes). After a 10-min incubation in the lysis buffer, an additional 10 strokes were performed with a tight pestle. The lysis procedure was monitored by light microscopy to ensure complete tissue and cell lysis and efficient nuclear extraction. The crude lysate was successively passed through 100 and 30µm mesh filters. The final lysate was spun at 1000 × g for 5 min and the resulting pellet was resuspended in 500 µl staining buffer (5% BSA in PBS) supplemented with RNAse inhibitor. Isolated nuclei were incubated with rabbit polyclonal antibodies specific for pericentriolar material 1 (PCM1) (Sigma-Aldrich; HPA023370) at a dilution of 1:400 for 1 h rotating at 4 °C. Next, Alexa Fluor 647-conjugated donkey-anti-rabbit 647 antibodies (ThermoFisher Scientific A-31573; 1:500 dilution), and DAPI (1:1000 dilution) were added and the incubation was continued for an additional hour. Samples were spun at 1000 × g for 10 min and washed with 500 µl staining buffer before resuspension in 500 µl staining buffer supplemented with RNAse inhibitor. Intact cardiomyocyte nuclei were sorted on a BD Influx FACS on the basis of DAPI and Alexa Fluor 647 positivity into cold BL+TG buffer from the ReliaPrep RNA Tissue Miniprep System (Promega, Z6112) for RNA isolation. RNA was isolated following a gDNA depletion step according to the manufacturer’s instructions.

### RNAseq

For RNA isolated from cardiac nuclei, 500 pg was used for library generation with the Ovation RNA-seq v2 kit (NuGEN M01206) kit. Libraries were prepared with the UltraLow V2 (NuGen) kit and sequenced on the HiSeq4000 system (Illumina) with 50 bp single-end reads. 4 control and 4 *Wt1^+/-^* samples were sequenced. Reads were mapped to the mm10 build of the mouse transcriptome using STAR^61^. Differential expression analysis was performed using the DESeq2 package based on a negative binomial distribution model. P-values were corrected for multiple testing using the false discovery rate (FDR) method of Benjamini-Hochberg. We have used 0.05 as FDR control level. PANTHER32 was used for gene ontology (GO) biological process analysis. Benjamini–Hochberg correction was performed for multiple testing-controlled P values. Statistically significant enriched terms were functionally grouped and visualized.

### RTqPCR

Total RNA was isolated from (sub) epicardial and (sub) endocardial ventricular sides of 12-16-(*Wt1+/-*) or 6-week-old (*Wt1BAR/BAR*) male and female mice using ReliaPrep RNA Tissue Miniprep System (Promega, Z6112) according to the manufacturer’s protocol. cDNA was reverse transcribed with oligo dT primers from 500ng of total RNA, according to the manufacturer’s protocol of the Superscript II Reverse Transcriptase system (Thermo Fisher Scientific, 18064014). Expression levels of candidate target genes were determined by quantitative real-time PCR using a LightCycler 480 Instrument II (Roche Life Science, 05015243001). Expression levels were measured using LightCycler 480 SYBR Green I Master (Roche, 04887352001) and the primers had a concentration of 10 pmol/L. The amplification protocol consisted of 5 minutes 95 °C followed by 45 cycles of 10 seconds 95°C, 20 seconds 60°C and 20 seconds 72°C. Relative start concentration (N0) was calculated using LinRegPCR^62^. Values were normalized to one or two reference genes per experiment (Hprt and Ppia). The primer sequences are as follows: (all 3’ to 5’) Hprt: TGTTGGATATGCCCTTGACT, GATTCAACTTGCGCTCATCT; Ppia: GGGTGGTGACTTTACACGCC, CTTGCCATCCAGCCATTCAG; Scn5a: GGGACTCATTGCCTACATGA, GCACTGGGAGGTTATCACTG; Wt1: TTCAAGGACTGCGAGAGAAGG, TATGAGTCCTGGTGTGGGTCT.

### *In vivo* electrocardiogram

12-16 (adult) or 65-67 week old male mice were anesthetized with 5% Isoflurane (Pharmachemie B.V. 061756) and placed on thermostated mat (36°C) with a steady flow of 1.5% isoflurane during all experiments. Electrodes were inserted subcutaneously in the limbs and connected to an ECG amplifier (Powerlab 26T, AD Instruments). The electrocardiogram (ECG) was measured for 5 minutes. ECG parameters were determined in Lead I based on 30 seconds of the recording. Only male mice were used given that BrS is more prevalent in males.

### Optical mapping and ajmaline exposure

Following ECG measurement, mice were sacrificed by cervical dislocation. The hearts were excised, cannulated and mounted on a Langendorff perfusion setup. Hearts were submerged in, and retrogradely perfused with 37 °C Tyrode’s solution containing (in mmol/L): NaCl 128, KCl 4.7, CaCl_2_ 1.45, MgCl_2_ 0.6, NaHCO_3_ 27, NaH_2_PO_4_ 0.4 and glucose 11 (pH maintained at 7.4 by equilibration with a mixture of 95% O_2_ and 5% CO_2_). The excitation-contraction uncoupler blebbistatin (10 μmol/L, Bio-Techne Ltd) was added in order to prevent movement artifacts. During the optical mapping experiments, pseudo-electrograms were recorded by placing three electrodes at ∼0.5 cm distance of the heart in the Einthoven configuration. After ∼10 minutes recovery period, hearts received a 0.4 ml bolus of 20 μmol/L Di-4 ANEPPS (Molecular Probes), the dye used for visualizing optical action potentials. Only hearts showing a regular sinus rhythm that was stabilized within the first two minutes were included for analyses. Conduction velocities were measured while pacing at the RV and the RVOT at a cycle length of 120 ms at twice the stimulation threshold. Hereafter, hearts were treated continuously with ajmaline (1.5 μmol/L, Carinopharm) and after a 10 minutes incubation period, the protocol was repeated. The average length of the full protocol was 66.7± 1.1 minutes. Optical signals were analyzed with previously published software ^63^, using Matlab2018. In short, optical signals were spatially (7×7 bin) and temporally (0-150 Hz) filtered and local activation was defined as the maximum positive dV/dt of the depolarization phase of the action potential. Conduction velocity was analyzed in two transversal-(RV and RVOT) and two longitudinal (RV) directions and averaged for each direction.

### Generation, dissociation and transfection of hiPSC-CMs

The hiPSCs (LUMC0099iCTRL04 line) were maintained on growth factor reduced matrigel (Corning, USA)-coated plates in mTeSR1 (Stem Cell Technologies, Canada) medium at 37°C (with 5% CO_2_ and 20% O_2_) and passaged every 3-4 days.

Cardiomyocytes were derived by using an adaptation of a published method^45^. Briefly, upon reaching 85-95% confluence differentiation of the hiPSCs started by switching the medium to RPMI 1640 (ThermoFisher Scientific, USA) supplemented with B27 minus insulin (ThermoFisher Scientific, USA) and 213ug/ml L-Ascorbic Acid 2-phosphate (hereunder referred to as-I+AA medium) and containing 6 µM CHIR99021 (Selleckchem, USA) for 72 hours. The medium was than replaced with-I+AA medium containing 2 µM Wnt-C59 (MedChemExpress, USA) for Wnt signaling inhibition and kept for other 72 hours on the cells. From day 8 onward, hiPSC-CMs were cultured in RPMI 1640 supplemented with B27 plus insulin (ThermoFisher Scientific, USA), without ascorbic acid and refreshed every 3-4 days until day 30.

For transfection and subsequent single-cell electrophysiological analysis, hiPSC-CMs cultures were dissociated with 1× TrypLE Select Enzyme (ThermoFisher Scientific, USA) for 10 minutes at 37⁰C, collected, centrifuged and reseeded at a density of approximately 4×10^3^ cells per coverslip in presence of RPMI 1640 supplemented with B27 plus insulin and L-Ascorbic Acid 2-phosphate (+I+AA medium). The glass coverslips were precoated with GFR matrigel/gelatin at 50/50 ratio.

Transfection was performed using Lipofectamine 3000 reagent (ThermoFisher) according to manufacturer’s protocol. Two pcDNA3.1 plasmids with CMV promotor and a P2A-eGFP site, containing one of two common *WT1* isoforms were used together in 1:1 ratio: isoform D (NM_024426.4, OHu27351, Genscript) and isoform B (NM_024424.3, OHu27419, Genscript). For the control, the same plasmid without *WT1* was used (“GFP-only”). Amount of plasmid (in ng) was chosen based on plasmid size (WT1-GFP 7.7kb, GFP 6.1kb), to ensure equal plasmid copies per condition. For each well in a 24-well plate, 0.75µl (for WT1) or 0.59µL (for GFP-only) lipofectamine was dissolved in 25µl RPMI (without glucose) and 500ng WT1 or 396 ng GFP-only plasmid with 1µl (for WT1) or 0.79µl (for GFP) P3000 in 25µl RPMI, after which these two were gently mixed (i.e. lipofectamine µl to DNA µg ratio 1.5) and incubated 10-15 minutes. Mixes were added to the cells and removed and replaced by culture medium after 3 days, and patched the next day.

### RV cardiomyocyte isolation

Freshly excised hearts of male mice were cannulated and mounted on a Langendorff setup, retrogradely perfusing oxygenated modified Tyrode’s solution at 37 °C containing (in mmol/L): NaCl 140, KCl 5.4, CaCl_2_ 1.8, MgCl_2_ 1.0, glucose 5.5, and HEPES 5.0; pH 7.4 (adjusted using NaOH). After 10 minutes, hearts were perfused with nominally calcium-free solution containing (in mmol/L): CaCl_2_ 0.01, creatine 10.7. After a consecutive 10 minutes, hearts were digested by addition of Liberase TM (0.032 mg/mL, Roche) and elastase (1.6 U/mL, SERVA Electrophoresis GmbH), while perfusing for ∼15 minutes. Hearts were washed in calcium-free solution containing 1% bovine serum albumin fraction V, fatty acid free (Roche). The RV was removed and cut into small pieces after further dissociating by pipette trituration to obtain single cardiomyocytes, which were washed in perfusion solution while up-titrating CaCl_2_ to 1.8 mmol/L. Isolated RV cardiomyocytes were stored at room temperature in modified Tyrode’s solution and used for AP and I_Na_ recordings within 2-3 hours.

### Cellular electrophysiology

*Data acquisition and analysis:* APs and I_to1_ were recorded with the amphotericin-B perforated patch-clamp technique while I_Na_ was recorded with ruptured patch-clamp technique, using an Axopatch 200B patch-clamp amplifier (Molecular Devices, San Jose, CA, USA). Voltage control, data acquisition, and analysis were done using custom software^64^ and pClamp10.6/Clampfit (Molecular Devices, San Jose, CA, USA). Borosilicate glass patch pipettes (GC100F-10; Harvard Apparatus, Waterbeach, UK) with a tip resistance of 2-2.5 MΩ were used. Signals were low-pass-filtered with a cutoff of 5 kHz and digitized at 20 kHz for I_Na_, 30 kHz for I_to1_, and 40 kHz for APs. Cell membrane capacitance was calculated by dividing the time constant of the decay of the capacitive transient after a −5 mV voltage step from −40 mV by the series resistance^46^. Series resistance was compensated for >80%. I_to1_ and APs were corrected for the calculated liquid junction potential. Following the procedure of Veerman et al.^65^, we selected single, spontaneously beating, GFP^+^ or WT1 overexpressing hiPSC-CMS showing regular, synchronous contractions in modified Tyrode’s solution, after which this extracellular solution was switched to a solution suitable for specific measurements as indicated below. For mouse, recordings were made from quiescent cardiomyocytes.

*Action potential measurement:* APs were measured in hiPSC-cardiomyocytes and mouse cardiomyocytes at 37°C in modified Tyrode’s solution, containing (in mmol/L): NaCl 140, KCl 5.4, CaCl_2_ 1.8, MgCl_2_ 1.0, glucose 5.5, HEPES 5.0; pH 7.4 (NaOH). Patch pipettes were filled with a solution containing (in mmol/L): K-gluconate 125, KCl 20, NaCl 5 (hiPSC-CMs) or 10 (mouse), amphotericin-B 0.44, HEPES 10; pH of 7.2 (KOH). APs were elicited at 4 Hz in mouse cardiomyocytes and at 1 Hz in hiPSC-CMs by 3 ms, 1.2-1.4 x threshold current pulses through the patch pipette. To overcome the lack of the inward rectifying potassium current (I_K1_) in hiPSC-cardiomyocytes, which limits the functional availability of I_Na_ and I_to_^66^, we injected an 2 pA/pF in silico I_K1_ with kinetics of Kir2.1 channels through dynamic clamp, as previously described and validated^67^. AP parameters were analyzed from 10 consecutive APs and averaged.

*Sodium current measurement:* I_Na_ was measured at room temperature (∼20°C). The bath solution for I_Na_ recordings in mouse cardiomyocytes contained (in mmol/L): NaCl 7.0, CsCl 133, CaCl_2_ 1.8, MgCl_2_ 1.2, glucose 11.0, nifedipine 0.005, HEPES 5.0; pH 7.4 (adjusted with CsOH), while hiPSC-CMs were perfused with an external solution containing (in mmol/L): NaCl 20, CsCl 120, CaCl_2_ 1.8, MgCl_2_ 1.0, glucose 5.5, nifedipine 0.010, HEPES 5.0; pH 7.4 (adjusted with CsOH). Pipette solution was composed of (mmol/L): NaCl 3.0, CsCl 133, MgCl_2_ 2.0, Na_2_ATP 2.0, TEACl 2.0, EGTA 10, HEPES 5.0; pH 7.2 (adjusted with CsOH). I_Na_ was recorded with a double-pulse protocol from a holding potential of –120 mV (see inset of Supplemental Figure 2C and Figure 6A; cycle length of 5 seconds). During the first 500 ms depolarizing pulses, I_Na_ activates and the currents analyzed here are used to determine I-V relationships and the voltage dependency of activation. For the latter, the I-V relationships were corrected for driving force and normalized to maximal current. The second pulse is used to determine the voltage dependency of inactivation and currents were normalized to the largest I_Na_. I_Na_ was defined as the difference between peak and steady-state current and densities were calculated by dividing current amplitude by C_m_. Voltage dependency curves of activation and inactivation were fitted with Boltzmann function: I/I_max_=A/{1.0+exp[(V_1/2_-V)/k]}, where V_1/2_ is the half-maximal voltage of (in)activation, and k is the slope factor (in mV). The time course of current inactivation was fitted by a double-exponential equation: I/I_max_=A_f_ ×exp(-t/τ_f_)+A_s_×exp(-t/τ_s_), where A_f_ and A_s_ are the fractions of the fast and slow inactivation components, and τ_f_ and τ_s_ are the time constants of the fast and slow inactivating components, respectively.

*Transient outward K^+^ current measurements.* I_to1_ was measured under conditions similar as mentioned for APs, but 0.25 mmol/L CdCl_2_ was added to the modified Tyrode’s to blocks I ^68^ which also strongly inhibits I ^69^. Suppression of these inward currents allows a more accurate determination of I_to1_. I_to1_ was measured using a double pulse protocol from a holding potential of-80 mV (see inset Figure 6D; cycle length of 5 seconds). The first pulse was used to determine the I-V relationship; the second pulse was used to analyze the voltage dependency of inactivation. I_to1_ was defined as the difference between peak and steady state current and densities were calculated by dividing current amplitude by C_m_. Voltage dependence of inactivation curves was fitted with the above-mentioned Boltzmann function and time course of current inactivation was fitted by a mono-exponential equation: I/I_max_=A_f_ ×exp(-t/τ_f_).

### Analysis of human GTEx and snRNAseq data

The Genotype-Tissue Expression (GTEx) data were obtained from left ventricle *Transcripts Per Million* (TPM) on 2022-06-06 v10. Samples were enriched in cardiomyocytes using the *Gene Set Variation Analysis* (GSVA) tool^70^ (RRID:SCR_021058). The heatmap was generated from centered and scaled data, and a Pearson correlation was performed on TPM values, with p-values adjusted for multiple hypothesis testing (number of tests > 2) using *Benjamini-Hochberg correction*.

For the correlation between *WT1* and *SCN5A* in single-cell RNA-seq data from hiPSC-CMs, we used the dataset published by Canac *et al.* (2022)^71^. Cells were differentiated following the *Matrigel sandwich method*^72^ and sequenced at day 30 using the *Multi Tissue dissociation kit 3* (Miltenyi Biotec) in duplicate. For these analyses, cells were filtered with a minimum of 250 features and a maximum of 8200, based on the sample distribution, as well as a maximum of 7% mitochondrial gene. Doublets were removed using *DoubletFinder*^73^ (RRID:SCR_018771). Data were then combined into a single SeuratV5 object^74^ (RRID:SCR_016341). Normalization with regression of cell cycle, number of features, and mitochondrial percentage was performed using *SCTransform*^75^ (RRID:SCR_022146). Cells were subsequently integrated using the *RPCA* method and annotated via *Azimuth* from SeuratV5 package with reference of human embryonic heart cell atlas at 7 PCW published by Asp *et al.* (2019)^76^. Correlation analysis was performed similarly as for the GTEx data, using the Seurat data matrix.

To assess the relationship between *SCN5A* and *WT1* expression in human heart tissue, we leveraged the Reichart et al. dataset. Reichart et al. performed snRNAseq on 61 explanted end-stage cardiomyopathy hearts, as well as 18 hearts from non-failing donors. The authors performed snRNAseq across various ventricular regions, including the left ventricle, the right ventricle, the interventricular septum, and the apex. For our analyses, we restricted to the 18 non-failing hearts and removed nuclei with ‘unknown’ annotations, leaving a total of 266,219 high-quality nuclei. For the regression analyses, we performed pseudo-bulking on the count matrix, where counts for each gene were collapsed per donor/region combination. Lowly-expressed genes were removed using the function *filterByExpr()* from the *edgeR* package (using design: ∼0 + region + sex + development_stage), after which *SCN5A* and *WT1* remained in the dataset. Count data were normalized using the *DESeq2* package, using functions *DESeqDataSetFromMatrix()*, *estimateSizeFactors()*, and finally *counts()* with flag ‘normalized=TRUE’. To regress the normalized expression of *WT1* on the normalized expression of *SCN5A*, we utilized a mixed negative binomial regression model, implemented in R-package *glmmTMB*, with the formula: *WT1* ∼ *SCN5A* + sex + development_stage + region + (1 | donor).

### Histology

Human RV, LV and interventricular septum 7 μm histological sections were obtained from the department of pathology at Amsterdam UMC. Sections were stained with a WT1-antibody (Abcam/Epitomics AC9115, clone EP122) diluted 1:50 using HIER (heat-induced epitope retrieval) (Tris-EDTA, pH 9.0) at 100°C. Sections were examined with a Leica imaged on a DSM5000 widefield microscope.

Adult mouse hearts were fixed overnight in 4% paraformaldehyde in PBS, dehydrated in a ascending ethanol series, paraffin embedded and sectioned at 8 µm. Sections were deparaffinized with xylene, rehydrated in descending ethanol series and treated with HIER (citrate-based antigen unmasking solution, Vector, H-3300-250) at 100°C. After permeabilization (Triton-X 0.5%) and blocking (BSA 5%), sections were stained with anti-cTnT (1:400, HyTest Ltd 4T19/2), anti-GFP (1:300, Abcam Cat ab13970), anti-WGA-555 (1:200, ThermoFisher W32464) and DAPI. Stained sections were imaged and photographed with a Leica DM6 microscope.

### Flow cytometry

Cell isolations for flow cytometry experiments were carried out according to a previously published protocol^77^. In short, after extracting the heart, the ascending aorta was clamped and a collagenase buffer was injected into the apex of the left ventricle. Then the atria were removed and the right ventricular free wall, left ventricular free wall and septum were separately dissociated by trituration. After suspending cells in stop solution (containing FBS), they were passed a 100µm strainer, centrifuged at low speed and stained for the cardiac marker cTnT. For the latter, cells were incubated 15 minutes in permeabilizing-blocking buffer (PBS with 0.1% saponin and 5% FBS), incubated with anti-cTnT (HyTest Ltd Cat 4T19/2, 1:400) for 1 hour and with secondary antibody (AF647 donkey anti-goat IgG H+L, Life A21447, 1:200) for 15 minutes, all at room temperature. Cells were confirmed to be all cardiomyocytes using a negative control without primary antibody. For compensation control, littermates without Cre were used. Cells were dissolved in FACS buffer (PBS with 0.5% BSA and 2mM EDTA) and measured with a LSRFortessa Flow Cytometer (BD Biosciences). FlowJo was used for analysis. Forward and side scatter information was used for gating of non-debris single cells and from these cells, the amount of endogenous tdT positive and GFP positive cells was quantified. For each experiment *Scn5a*^+/+^ and *Scn5a*^+/-^ hearts were used in parallel, with researchers being blinded for the genotype during experiment and analysis.

### Statistics

The experimenters were blind to genotypes during all measurements and outcome assessment. Datasets were tested for normality using Shapiro-Wilk test unless specified otherwise. Whole isolated tissue RT-qPCR were analyzed with ANOVA with Welch’s correction followed by unpaired t-tests with Welch’s correction (Figure 1), or 2-way ANOVA followed by Šídák’s multiple comparisons test and adjusted p-values are shown (Figure 5). *In vivo* electrophysiology and optical mapping measurements were analyzed with unpaired t tests for 2 group comparisons, and 2-way ANOVA followed by Tukey’s multiple comparison test for aging experiments comparing 3 groups. For single cell electrophysiology, data are expressed as mean±SEM. Statistical analysis was carried out with SigmaStat 3.5 software (Systat Software, Inc., San Jose, CA, USA). Normality and equal variance assumptions were tested with the Kolmogorov–Smirnov and the Levene median test, respectively. Significance between parameters of two groups was tested using unpaired Student’s t-test, or in case of a failed normality and/or equal variance test, by Mann-Whitney Rank Sum Test. Data of I-V relationships was tested for significance using the two-way repeated measures ANOVA followed by pairwise comparison using the Student– Newman–Keuls test. P<0.05 defines statistical significance.

For the flow cytometry data, a general linear mixed model was used after rank transformation of GFP percentages due to non-normally distributed residuals. Bonferroni post-hoc pairwise comparisons of regions and Mann-whitney U pairwaise tests of genotypes were used.

Data are presented as individual data points and mean or mean ± standard error of the mean (SEM) and p<0.05 defines statistical significance. Statistical analysis was performed using GraphPad Prism 10 and SPSS.

## Acknowledgements

We acknowledge the resources and support from the Core Facility Genomics of the Amsterdam UMC for the RNA sequencing experiments. We are thankful to Jaël Copier, Caroline Pham, and Aya Aliawi for their technical assistance. We thank Vincent Christoffels for insightful discussions which were helpful in the advancement of this work.

## Supplemental Data

Supplemental figure legends

**Supplemental Figure 1.**
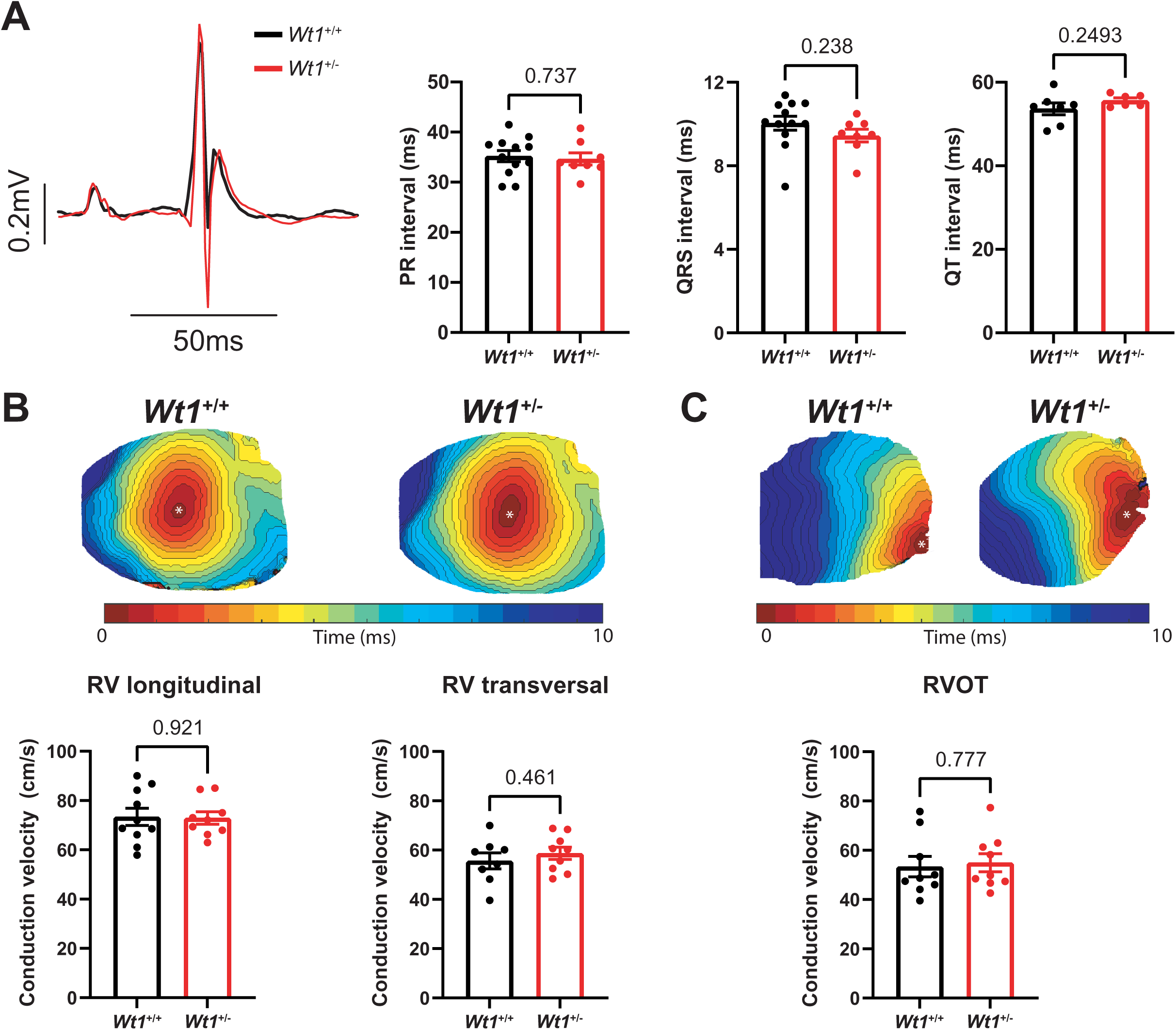
Heterozygous loss of *Wt1* (*Wt1^+/-^*) does not affect cardiac conduction in adult mice. A) Typical ECGs and average ECG parameters of control (n = 7-12) and *Wt1^+/-^* (n = 6-8) mice. B) Typical examples and average conduction velocities of longitudinal and transversal conduction in RV, and C) transversal conduction in RVOT in control (n = 9-10) and *Wt1^+/-^* (n = 9) mice. RV, right ventricle; RVOT, right ventricular outflow tract. Statistical significance was determined with unpaired t-tests.

**Supplemental Figure 2.**
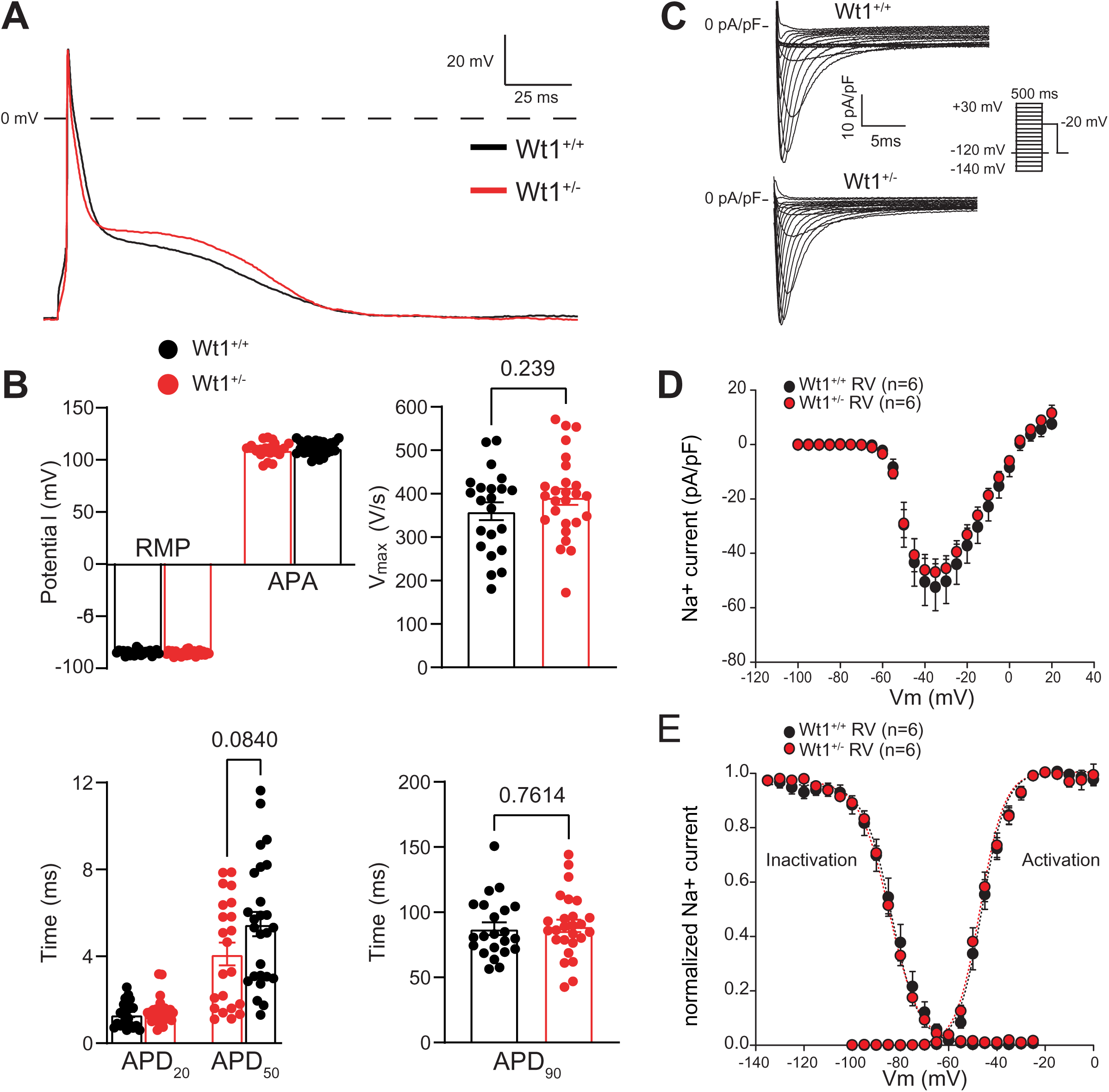
Heterozygous loss of *Wt1* does not affect electrical properties of single adult RV cardiomyocytes. A, B) Typical action potentials (AP; A) measured at 4 Hz and average AP characteristics (B) of control (n = 26) and *Wt1^+/-^*(n = 22) RV cardiomyocytes. C-E) Typical sodium currents (I_Na_; C), average I_Na_ current-voltage relationships (D), and voltage dependence of (in)activation (E) of control (n = 6) and *Wt1^+/-^* (n = 6) isolated RV cardiomyocytes. Dotted lines are Boltzmann fits. The inset in panel C shows the used voltage-clamp protocol to measure I_Na_. RMP, resting membrane potential; APA, AP amplitude; V_max_, maximal AP upstroke velocity; APD, AP duration; RV, right ventricle. Statistical significance was determined with two way RM ANOVA or unpaired t-tests.

**Supplemental Figure 3.**
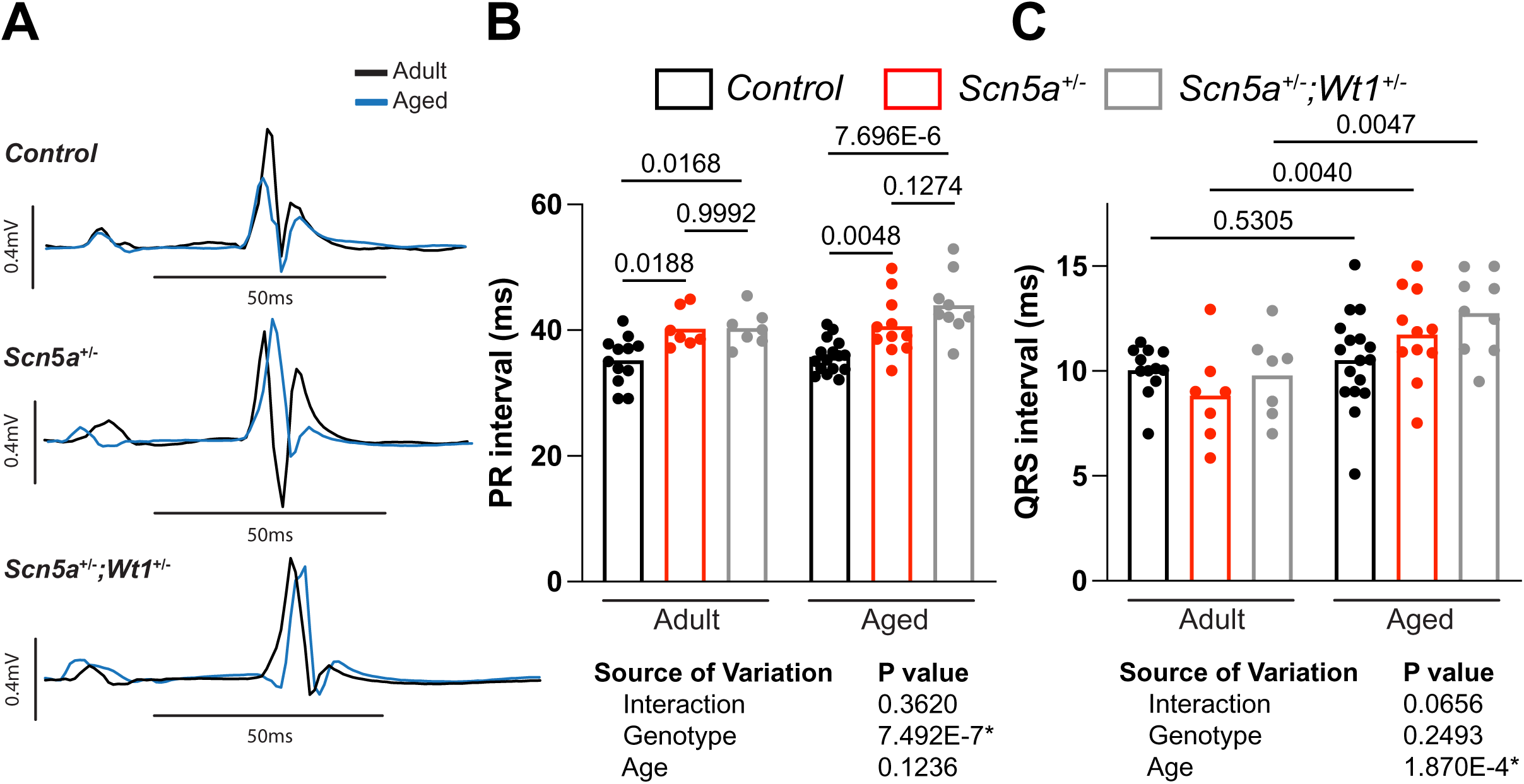
Effect of aging and diminished *Wt1* on ECG parameters of *Scn5a^+/-^* <u>mice.</u> A) Typical ECG traces and average duration of B) PR-and C) QRS duration of adult and aged control (n= 12 and 17, respectively) *Scn5a^+/-^* (n=7 and 11, respectively) and *Scn5a^+/-^*;*Wt1^+/-^*(n=7, 9 respectively) mice. Significance was determined using 2-way ANOVA shown in the tables below each set of graphs, followed by pairwise Tukey’s multiple comparisons tests (shown in graphs).

**Supplemental Figure 4.**
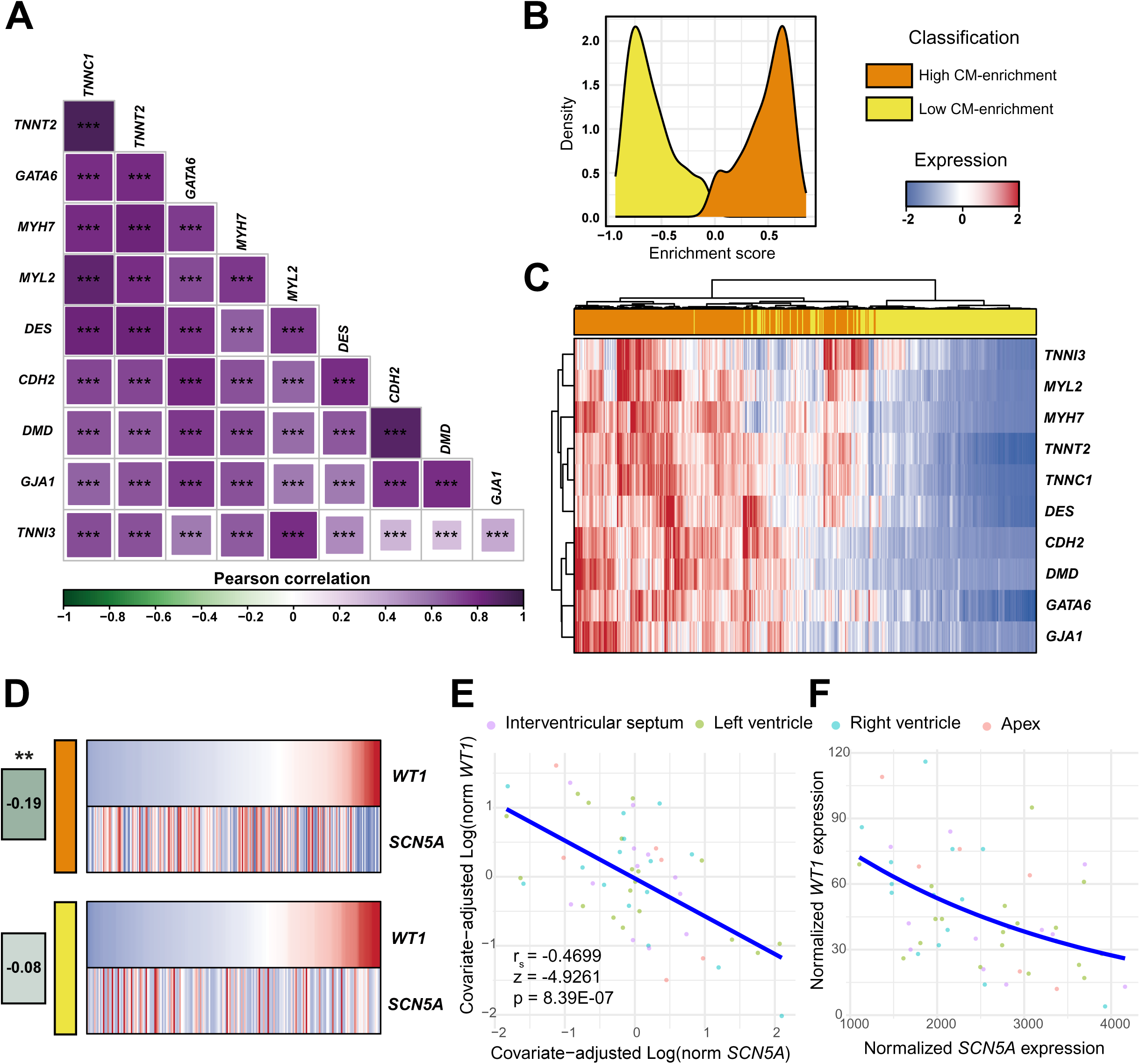
High *WT1* expression correlates with decreased *SCN5A* expression. A) Pearson correlation analysis of left ventricular samples from GTEx dataset using a selection of a CM 10-gene signature, as reported by^35^ (***: p-value<0.001). B) Distribution of GTEx sample based on their CM enrichment scores, using the 10-gene signature (Gene Set Variation Analysis method, GSVA R package). Samples were classified into “high” (orange) and “low” (yellow) CM-enrichment groups based on the bimodal distribution of scores. C) Heatmap displaying the expression values of left ventricular samples from GTEx dataset for each of the 10 genes that compose the CM signature. Blue and red indicate low and high expression values, as compared to the mean for each gene, respectively. Dendrogram was generated using Ward’s hierarchical clustering method with Euclidean distances. D) Pearson correlation of *SCN5A* and *WT1* expression for the “high” and the “low” CM-enrichment groups. E) Partial regression plot for mixed negative binomial model between *WT1* and *SCN5A* from 18 healthy donors across 4 ventricle regions in pseudo-bulk. Axes represent the covariate-adjusted log-transformed expression of *WT1* and *SCN5A.* Corresponding statistics are added, where r_s_ represents the Spearman’s correlation coefficient, while z and p represent the Z-value and P-value from the mixed negative binomial regression respectively. F) Univariate negative binomial model plot with untransformed data from E.

**Supplemental Figure 5.**
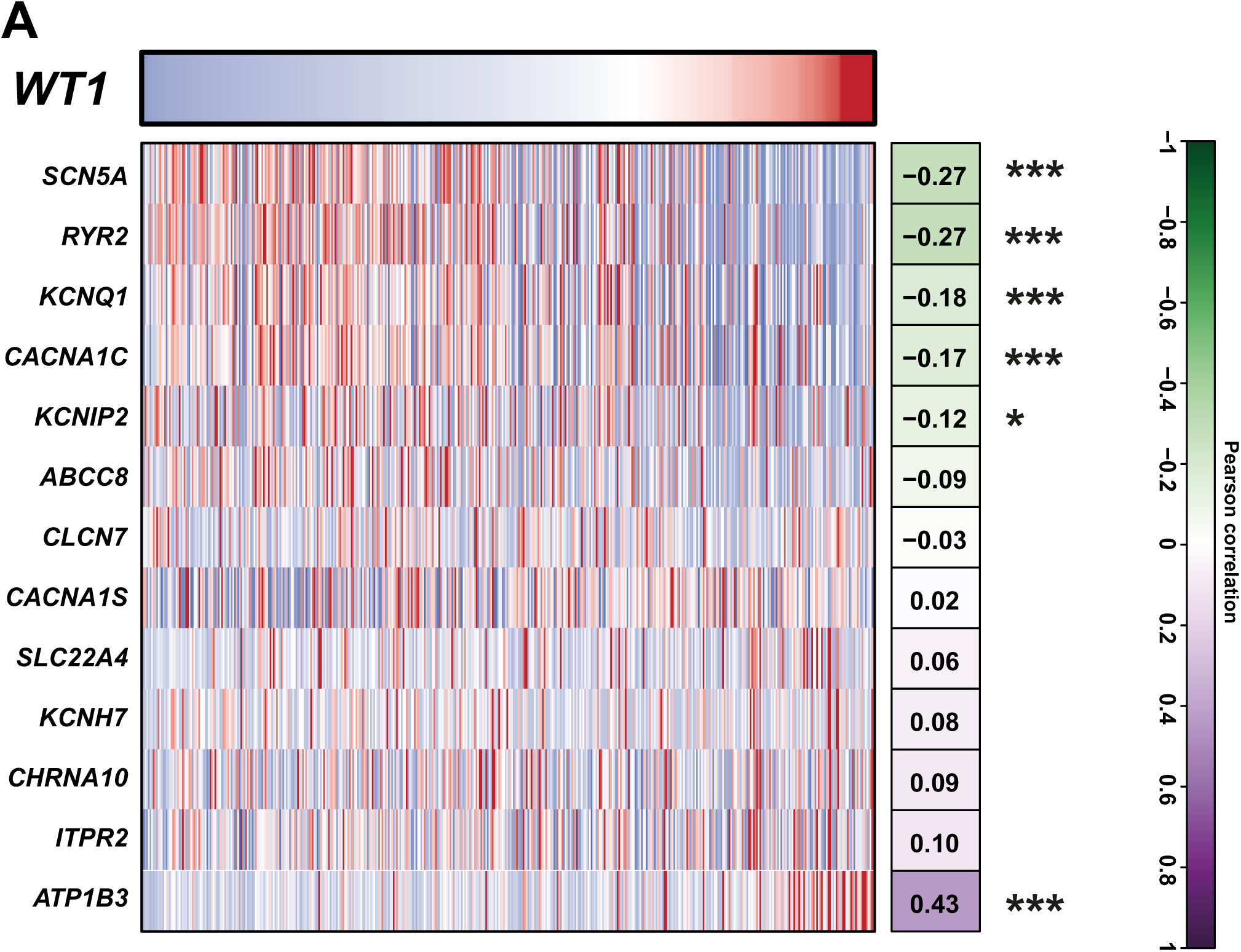
Conduction-related genes altered in cardiomyocytes of *Wt1^+/-^* mice, also correlate with *WT1* expression in human left ventricular samples. Pearson correlation analysis of *WT1* expression and of the conduction-related genes altered in cardiomyocytes of *Wt1^+/-^*mice (Figure 1D), using human left ventricular samples from GTEx database. *: p-value<0.05; ***: p-value<0.001. All p-values were adjusted for multiple hypothesis testing using Benjamini-Hochberg correction.

**Supplemental Figure 6.**
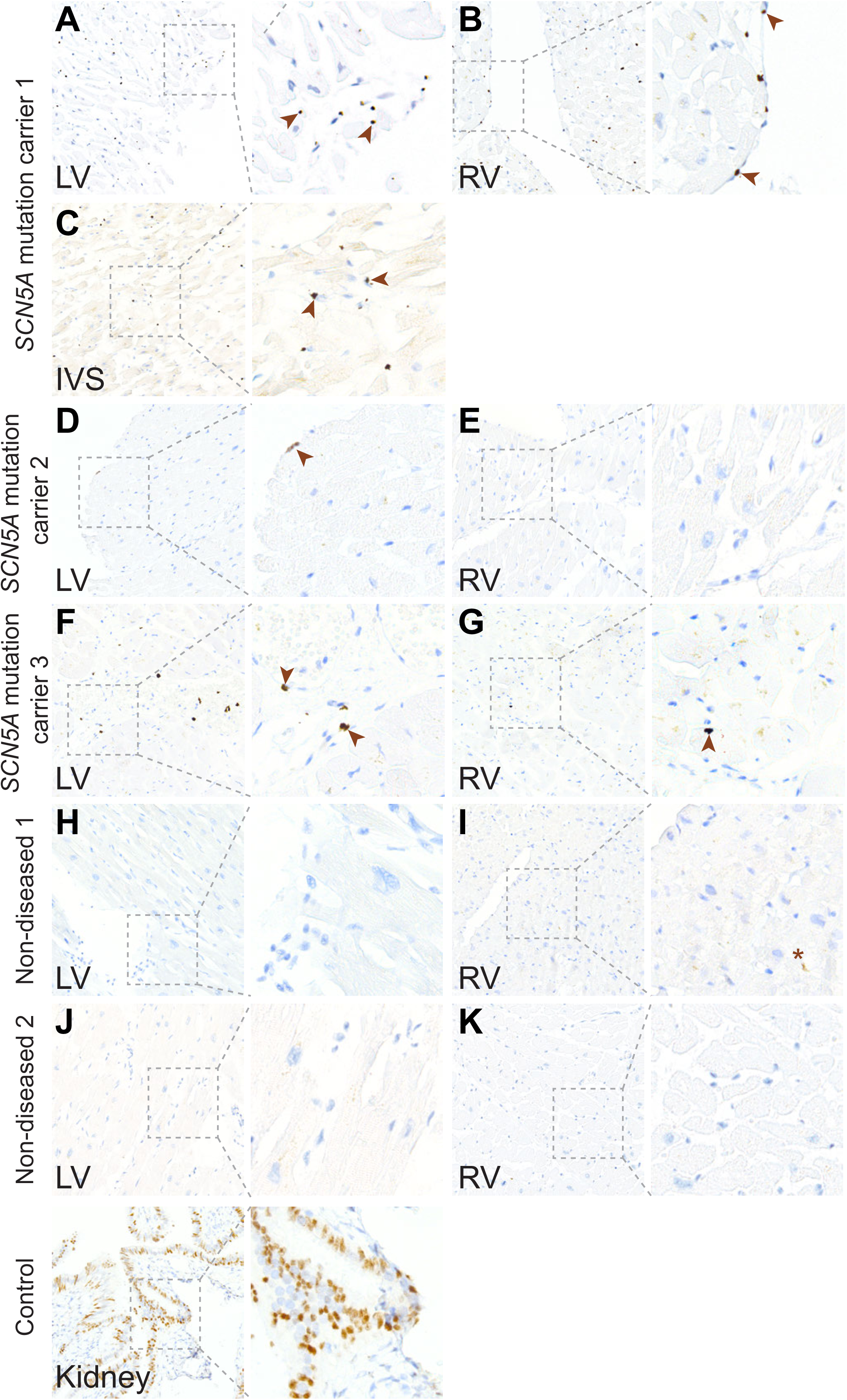
Immunohistochemistry for WT1 in cardiac sections of patients with *SCN5A* mutation and non-diseased hearts. A-K) Representative images of *SCN5A* mutation carriers (A-G) and non-diseased (H-K) left ventricles (LV (A, D, F, H, J), RVs (B, E, G, I, K), and interventricular septum (IVS) (C) histological sections. L) Staining control in human kidney. Arrowheads denote nuclear staining. Asterisk denotes non-specific signal.

---

**Supplemental Table 1.** DEseq2 analysis of control vs *Wt1^+/-^* cardiomyocytes.

**Supplemental Table 2.** Average sodium current (I_Na_) characteristics of cardiomyocytes isolated from the right ventricle of control and *Wt1^+/-^* mice.

| Sodium Current |  |  |  |
| --- | --- | --- | --- |
| Parameter | Control (n=6) | *Wt1*<sup>+/-</sup>(n=6) | P value |
| Maximal peak $I_{Na}$ (pA/pF) | -52.5 ± 8.6 | -46.9± 4.8 | 0.58 |
| $V_{1/2}$ activation (mV) | -45.55 ± 1.26 | -46.25 ± 1.28 | 0.706 |
| k activation (mV) | 4.91± 0.32 | 5.21± 0.31 | 0.519 |
| $V_{1/2}$ inactivation (mV) | -83.23 ± 1.98 | -84.21 ± 0.89 | 0.661 |
| k inactivation (mV) | -6.20± 0.32 | -6.11± 0.46 | 0.885 |

**Supplemental Table 3.** Region-specific *WT1-SCN5A* correlation analysis in all cell-types.

| Region | N donors | N cells | N donor region combinations | Spearman Rho | Beta | SE | Z | P |
| --- | --- | --- | --- | --- | --- | --- | --- | --- |
| All | 18 | 266219 | 51 | -0,4699548 | -4,93E-04 | 1,00E-04 | -4,9260962 | 8,39E-07 |
| IVS | 12 | 51299 | 12 | -0,3286713 | -5,22E-04 | 5,71E-04 | -0,9148981 | 3,60E-01 |
| RV | 15 | 58779 | 15 | -0,5214286 | -7,25E-04 | 3,86E-04 | -1,8766413 | 6,06E-02 |
| LV | 18 | 132742 | 18 | -0,246646 | -2,40E-04 | 7,16E-05 | -3,3486163 | 8,12E-04 |
| Apex | 6 | 23399 | 6 | -0,8857143 | -9,32E-04 | 1,78E-04 | -5,246449 | 1,55E-07 |

Model: expression\_gene1~expression\_gene2+gene+sex+age

**Supplemental Table 4.**
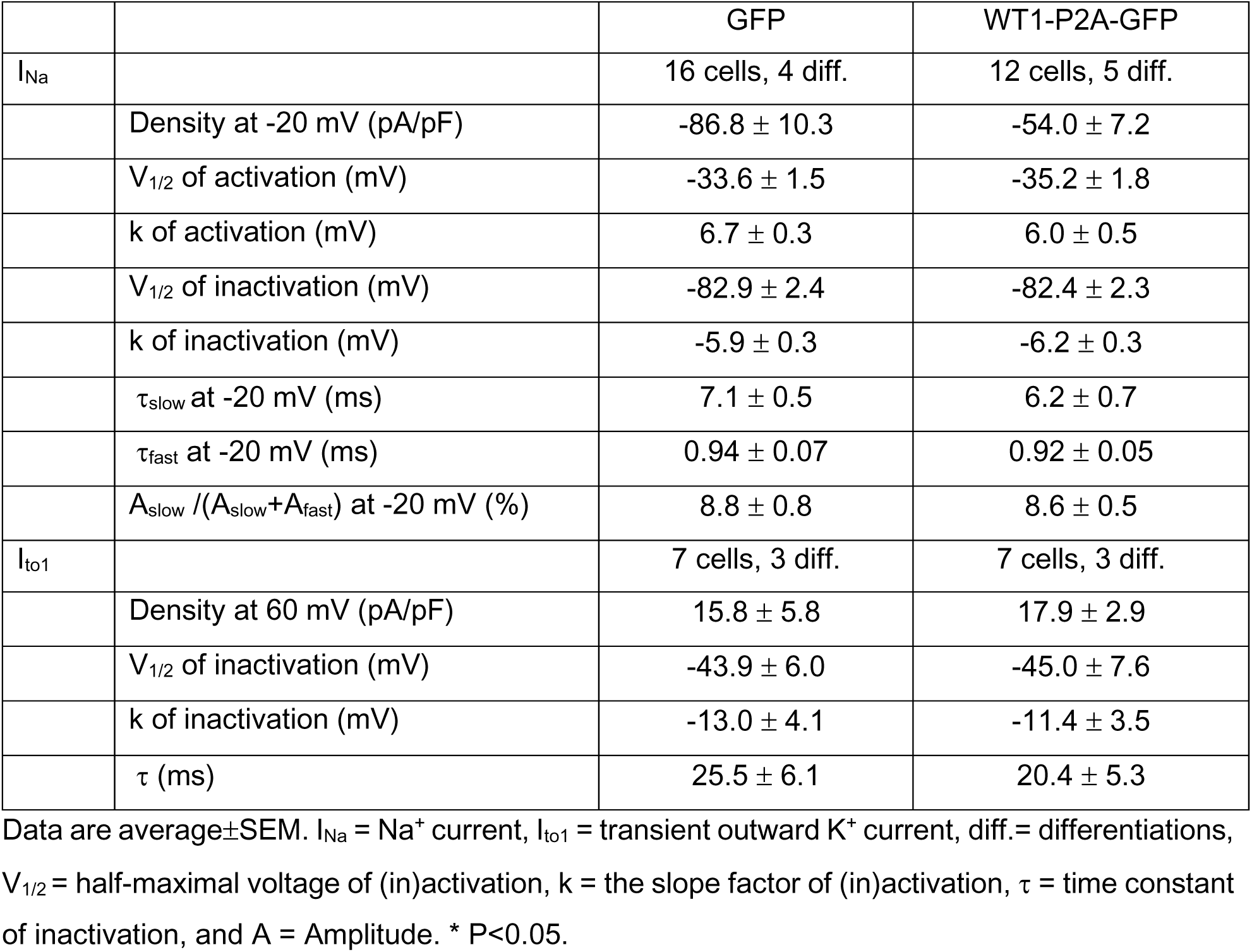
I_Na_ and I_to1_ parameters in hiPSC-cardiomyocytes.

